# New high accuracy diagnostics for avian *Aspergillus fumigatus* infection using Nanopore methylation sequencing of host cell-free DNA and machine learning prediction

**DOI:** 10.1101/2025.04.11.648151

**Authors:** Markus Hodal Drag, Christina Hvilsom, Louise Ladefoged Poulsen, Henrik Elvang Jensen, Stamatios Alan Tahas, Christoph Leineweber, Carolyn Cray, Mads Frost Bertelsen, Anders Miki Bojesen

## Abstract

Avian aspergillosis is a detrimental fungal infection affecting wild and domestic birds yet sensitive antemortem diagnostics for early clinical infections are lacking. Here we present new diagnostics for *Aspergillus fumigatus* (*Af*) infection developed from cell-free DNA (cfDNA) methylation markers. Broiler chickens were experimentally infected with either *Af,* a non-*Af* agent (*Escherichia coli or Gallibacterium anatis*) or assigned as controls. Oxford Nanopore (ONT) sequencing was performed on serum cfDNA (n = 124), and machine learning (ML) models were trained on infection-specific markers. Three tests were developed: A ‘High Accuracy’ test for best performance (sensitivity: 100%, specificity: 89.2%) and robustness (ROC-AUC: 0.92) as well as ‘Fast’- and ‘In situ’ tests for rapid turnaround and methylation PCR. Diagnostic accuracies were 92.3%, 82.7%, and 73.1%, respectively. In conclusion, new tests using on ML- and host cfDNA methylation markers demonstrated high diagnostic performance comparable to microbial cfDNA (mcfDNA) tests but without concern for environmental contamination.

**Key highlights:**

- We present three new high accuracy diagnostic tests for *Aspergillus fumigatus* infection in chickens that use methylation markers from serum cell-free DNA (cfDNA).
- Differentially methylated cfDNA regions (DMRs) were detected by Oxford Nanopore sequencing (ONT) in experimentally infected chickens and used as markers to train machine learning (ML) models for development of three diagnostic tests.
- The highest accuracy was found with 83 markers of 10 kilobases (KB) using the glmnet algorithm for the ML model, which classified 92.3% blinded samples correctly.
- A Fast test designed for cheap <1h sequencing using adaptive sampling could correctly classify 82.7% samples with 22 markers using a random forest (rf) model.
- An In situ test with only four markers, envisioned for use in a simple methylation-specific PCR (MSP-PCR) assay, could correctly classify 73.1% blinded samples.
- Reference values with associated probabilities of infection were calculated for each of the three tests and are presented for further evaluation.

## Introduction

Avian aspergillosis is a fungal respiratory infection caused by the ubiquitous mould *Aspergillus fumigatus* (*Af*) ^1^ affecting wild and domestic birds ^2–4^. In zoo-housed birds, notably penguins and birds of prey, aspergillosis is one of the most prominent causes of morbidity ^5–8^. In poultry production, the infection can lead to significant economic losses ^9^. Antemortem diagnosis is usually based on cumulative evidence from clinical observations, fungal culture, endoscopy, diagnostic imaging as well as haematology, serum protein electrophoresis (SPE), liquid chromatography mass spectrometry (LC-MS) of gliotoxin ^10^, serum amyloid A (SAA) levels ^11^, galactomannan and PCR using blood ^12^, and immunological blotting of antibodies against *Af* ^13^. However, no single diagnostic technique is characterised by high sensitivity (SE), specificity (SPE), cost-effectiveness, minimally invasiveness, and ability to detect infection well ahead of serious illness.

Circulating cell-free DNA (cfDNA) are short (∼167 bp) DNA fragments released from apoptotic or necrotic cells and subsequently expelled to blood, urine and other bodily fluids ^14^. As the analytical backbone of the “liquid biopsy” concept ^15–17^, cfDNA has been investigated as a biomarker for a wide variety of conditions and diseases in mammals: infection ^18^, mitochondrial ^19^ and metabolic disorders ^20^, obesity ^21^, inflammation ^22^, physical stress ^23^, impact of space flight ^24^, and aging ^25^. Strategies include assessing changes in concentrations and targeted approaches seeking to analyse genetic markers and chemical moieties in the cfDNA. However, concentrations alone can be difficult to interpret due to natural tissue turnover ^26–28^, thus requiring deeper qualitative interrogation such as backtracking to tissue of origin by nucleosome footprints ^29^, fragmentation patterns ^30^, and methylation patterns ^20,31–33^. For the latter, Oxford Nanopore Sequencing (ONT) can directly measure 5-hydroxymethylcytosine (5hmC) and 5-methylcytosine (5mC) ^34–40^, which can be used to identify tissue-specific differentially methylated regions (DMRs) ^20^. Previous studies have demonstrated tissue overcontribution of cfDNA during COVID-19 ^41^, as well as association between methylation and severity of infection ^42,43^, as well as increased cfDNA methylation in patients with active pulmonary tuberculosis (TB) compared to latent TB ^44^.

We hypothesised that *Af*-specific DMRs could be identified with ONT sequencing and used to train machine learning (ML) models for *Af* prediction. The aims were to i) create an *in vivo Af* model in chickens and successfully conduct ONT sequencing on cfDNA from 200 µl serum; ii) to find infection-significant DMRs and use for ML ^45^ to predict infection; iii) to evaluate by a performance cohort with chickens infected with *Af,* E*. coli* or *G. anatis* as well as naturally exposed chickens; and iv) to select three ML models as new diagnostic tests with the highest possible accuracies but with distinguishable capabilities with respect to turnaround speed and possibility to adapt to methylation PCR (Fig. 1).

**Fig. 1:**
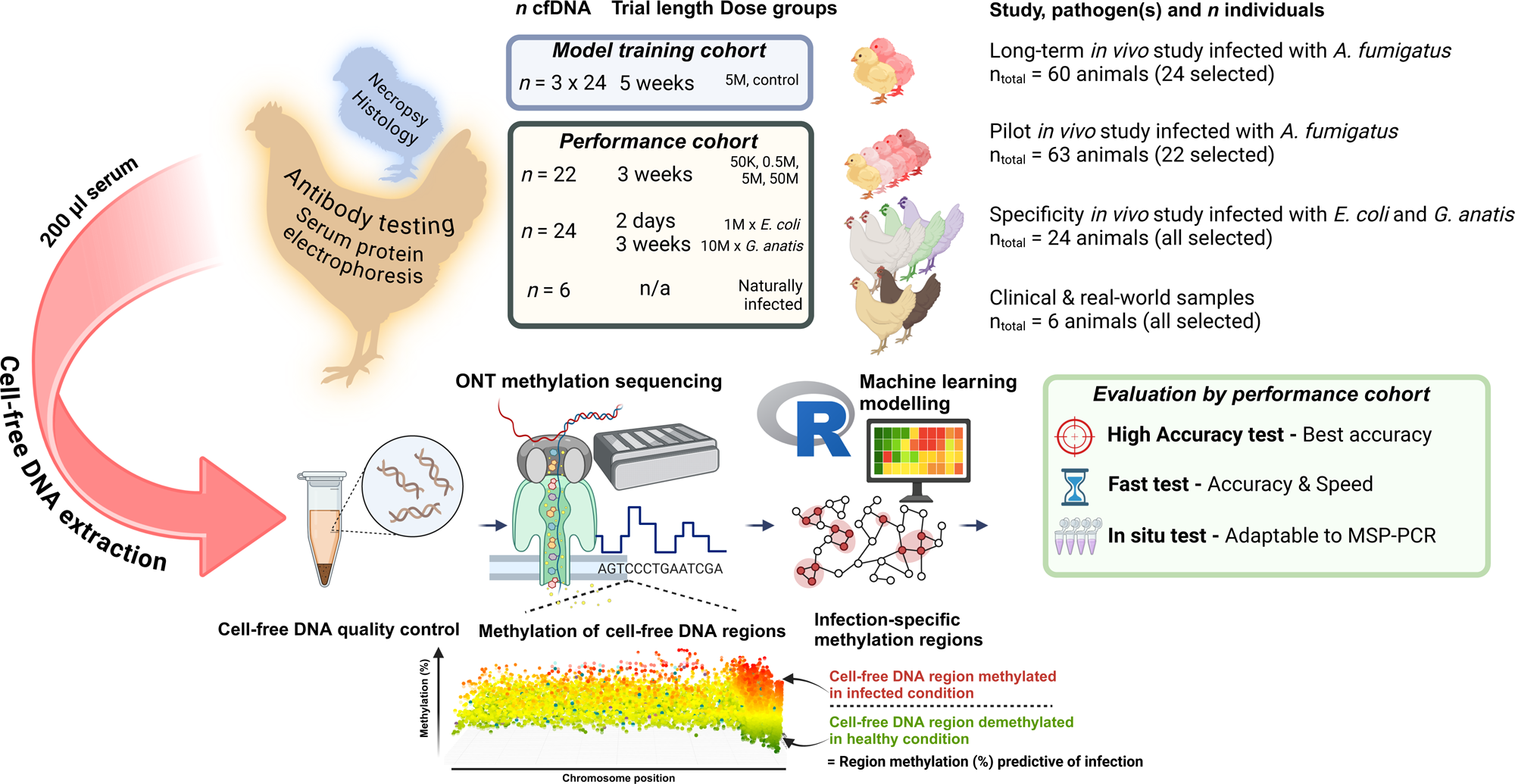
A total of 124 serum samples from a training (n = 72) and performance cohort (n = 52) were subjected to cell-free DNA extractions and sequenced with Oxford Nanopore methylation sequencing. Three tests were generated to serve different diagnostic requirements of accuracy, speed, and portability. Created with BioRender.com.

## Results

### Quality control of infection doses by semi-quantitative real-time PCR

Comparison between microscopy enumerations and conidial equivalents (CE) calculated from *FKS* gene copies showed median *FKS* copies (± IQR) of 5.04M (± 0.29M) and 4.89M (± 0.41M) for the two 5M groups in the pilot- and long-term study, respectively, which was statistically non-significant. The 0.5M dose was significantly (p = 0.05) higher in the long-term study but was not used in the *in vivo* study, only for quality control purposes (Fig. 2A). PCR efficiency was ≥ 91%. Full results and *FKS* sequence are available in Supplementary File 1. The CE *FKS* copies between pilot- and long-term study exhibited a significant correlation (Spearman rank-correlation *R* = 0.89, *p* = 0.0014), demonstrating that titrations over the three log^10^ steps exhibited high similarity (Fig. 2B). Finally, light microscopy confirmed the morphology of *Af* conidiophore for species validation (Fig. 2H).

**Fig. 2:**
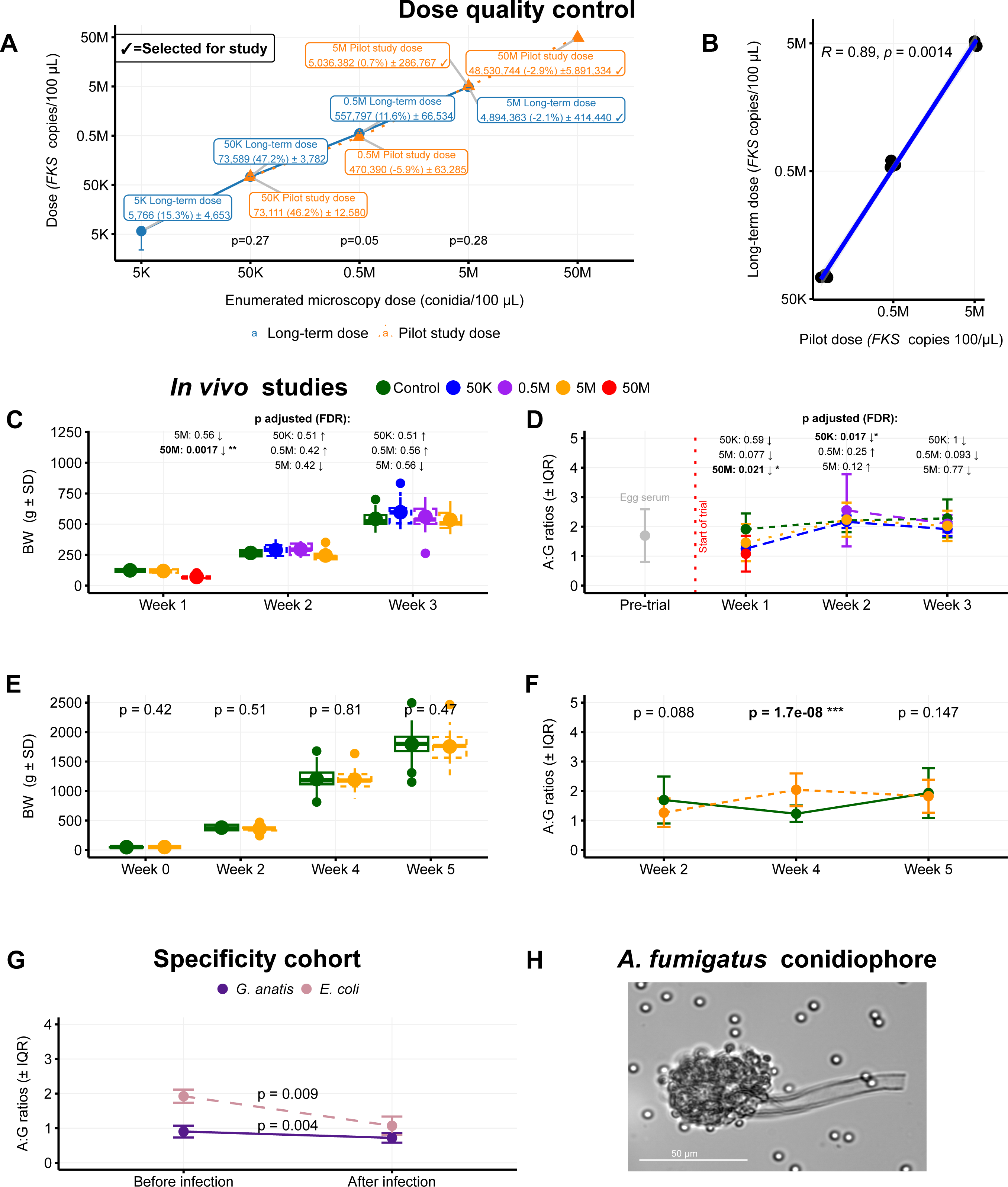
A) Microscopy compared with median (± IQR) *FKS* copies; B) Inter-study dose correlation; C,E) Weights for pilot-(n = 63) and long-term study (n = 60); D,F,G) Albumin:globulin (A:G) ratios for pilot-(n = 142), long-term-(n = 171) and specificity cohort (n = 24); H) Microscopy (400x) of conidiophore.

### *In vivo* studies for dose determination and model training

For the pilot study, a significant decrease (FDR < 0.05) in mean body weight was observed in the 50M group compared with controls at the first week of trial. No significant difference was observed in mean body weight after three weeks in any of the dose groups compared with controls (Fig. 2C). For the 50M group, euthanasia was performed by cervical dislocation during day 2 to day 4 due to grave and persistent symptoms of acute aspergillosis, including severe respiratory distress and dyspnoea, tachypnoea and silent gasping. The final mean body weight for the 50M group was 70.8 g (± 17.1 g), which was measured postmortem before necropsy and scoring. No significant differences were observed in mean body weights in the long-term study between 5M group and controls at any week (Fig. 2E) and no clinical signs of acute or chronic aspergillosis were observed.

### Serum protein electrophoresis – *in vivo* studies and specificity cohort

In the pilot study, median albumin:globulin (A:G) ratios decreased significantly (FDR < 0.05) for dose group 50M in the first week compared to the control samples. For week 2, A:G ratios decreased significantly (FDR < 0.05) for dose group 50K compared to controls, respectively (Fig. 2D). In the long-term study, median A:G ratios were significantly (*p* < 0.05) higher in the 5M group in week 4. No significant difference between 5M and control samples were detected in week 2 or week 5 (Fig. 2F). In both *G. anatis* and *E. coli* groups in the specificity cohort, A:G ratios dropped significantly (*p* < 0.05) after infection (Fig. 2G). The inter-assay variation expressed as the coefficient of variation (CV) was 7.3% and 17.9% for total protein (TP) and A:G ratios, respectively. Full SPE data for all tested samples (n = 342) comprising all animals in the *in vivo* studies (n = 123) across all time points of sampling is available in Supplementary file 2.

### Postmortem gross examinations

For the pilot study, statistical comparison by binomial test showed a total of 10 significantly different (FDR < 0.0001) scoring parameters across groups. These were found in the 50K and 0.5M groups compared with expected baseline values in airsac transparency, and all four infection dose groups were significantly different (FDR < 0.001) in the parameters of lung congestion, lung oedema, organ size, and granulomas (Fig. 3A).

**Fig. 3:**
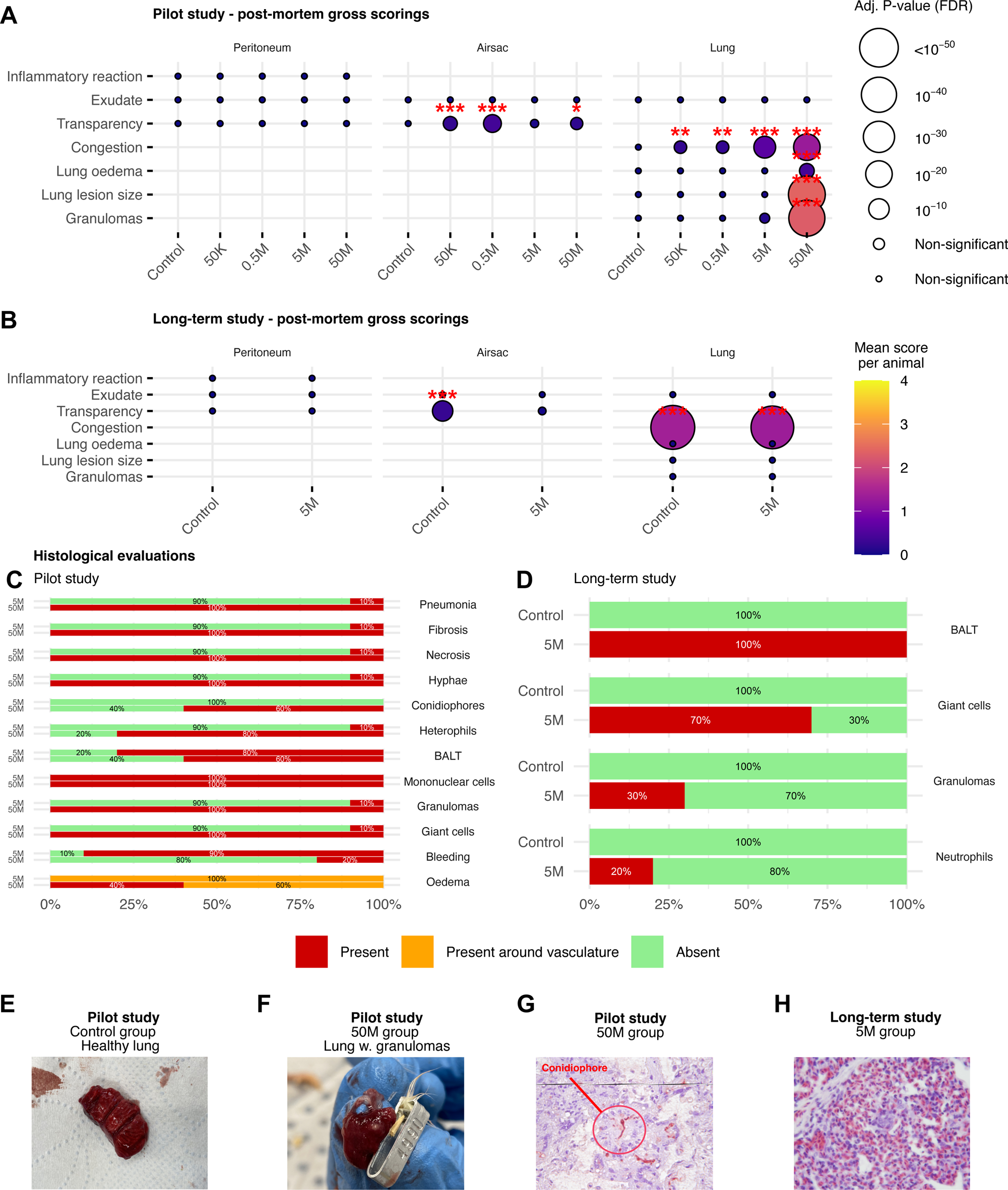
Postmortem for A) pilot study (n = 63) and B) long-term study (n = 60); C,D) histopathology of pilot (n = 15) and long-term study lungs (n = 20); E) healthy lung; F) granulomas; G) 50M group immunohistochemistry slide; H) fungal-negative H&E stained slide. BALT = Bronchus-Associated Lymphoid Tissue (BALT).

In the long-term study, airsac transparency scored significantly higher (FDR < 0.001) in the control group when compared against baseline values and for the lung congestion parameter, both control and 5M groups exhibited significantly different (FDR < 0.05) scorings (Fig. 3B). Only chickens from the 5M and 50M groups in the pilot study formed visible granulomas (Fig. 3F) compared with healthy control lungs from the same study (Fig. 3E). Full phenotype datasets with postmortem data for both *in vivo* studies are available in Supplementary file 3 and 4.

### Histopathology and immunohistochemistry

To confirm *Af* presence in the lungs, one left lung sample from the 50M group (ID: 4990) and one left lung sample from the 5M group (ID: 4983) from the pilot study were analysed using specific *Aspergillus* immunohistochemistry. For the 50M sample, the staining revealed the presence of multiple hyphae (Fig. 3G). For general histopathology, a significant increase (p < 0.05) in the frequencies of pathology (mean ± SD) was observed between 5M group (36.7% ± 41.6%) and 50M group (85% ± 25.8%) (Fig. 3C).

In the long-term study, no fungal remains were found by haematoxylin and eosin (H&E) staining and immunohistochemistry of six lung samples from the 5M group (IDs: 3430, 3432, 3433, 3437 3440 and 3448), as exemplified in Fig. 3H. A significant increase (p < 0.05) in the frequencies of pathology (mean ± SD) was found for the 5M group (55% ± 37%) compared to the controls, that were all negative (Fig. 3D).

### Serology-based diagnostics using chicken IgG

Of the 67 samples, no positive samples were identified among clinical, control or experimentally infected *in vivo* samples from the long-term or pilot studies for circulating *Af*-specific IgG antibodies (Table 1). Serum from a postmortem confirmed *Af-*infected penguin was found positive in every test as used as positive control. For all runs using the IgG test, sensitivity was 3.4%, specificity was 100%, and diagnostic accuracy was 64.6%. Full serology dataset is available in Supplementary file 5.

**Table 1:** Diagnostic evaluation of chicken IgG test targeted against *A. fumigatus* antigens.

| Cohort | *n* samples | TP | FP | TN | FN | Diagnostic accuracy |
| --- | --- | --- | --- | --- | --- | --- |
| Long-term study | 33 | 0 | 0 | 15 | 18 | 45.5% |
| Pilot study | 11 | 0 | 0 | 3 | 8 | 27.3% |
| Specificity cohort –<br>*E. coli* & *G. anatis* | 16 | 0 | 0 | 16 | 0 | 100% |
| Clinical & real-world samples | 6 | 0 | 0 | 3 | 3 | 50% |
| Positive penguin control | 1 | 1 | 0 | 0 | 0 | 100% |
| *Total* | 7 | 1 | 0 | 37 | 29 | 64.6% |

*n*: number; TP: true positive; FP: false positive; TN: true negative; FN: false negative.

### ONT sequencing of cell-free DNA from training and performance cohorts

For all extractions included in the study (n = 124), mean cfDNA (range) per sample was found to be 22.3 (7.4-57.5) ng/µl (Table 2 and Table 3). A total of 96 cfDNA samples were sequenced with the ONT PromethION P2 Solo device and the remaining 28 cfDNA samples were sequencing with the ONT MinION. For the long-term study, the mean sequencing reads per animal (± SD) were 10.1M (± 3.1M) and 9.6 (± 2.8) for control and 5M infected groups, respectively (Table 2). For the performance cohort (n = 52 cfDNA samples), mean sequencing reads per animal (±SD) were 3.4M (± 1.2M) and 3.9M (± 0.9M) for control and infected groups, respectively (Table 3).

**Table 2:** Mean cell-free DNA concentrations and sequencing reads obtained from the long-term study used for training machine learning models.

| Long-term study | | Avg. cell-free DNA (± SD)<br>(ng/μl per animal) | | Avg. reads (± SD)<br>(million reads/animal) | | ONT device |
| --- | --- | --- | --- | --- | --- | --- |
|  | *n* | Control | Infected | Control | Infected |  |
| Week 2 | 24 | 5.9<br>(± 4.1) | 6.3<br>(± 2.0) | 9.2M<br>(± 3.2M) | 10.2M<br>(± 1.8M) | P2 Solo |
| Week 4 | 24 | 12.8<br>(± 20.4) | 4.9<br>(± 2.9) | 13.1M<br>(± 3.1M) | 8.3M<br>(± 3.6M) | P2 Solo |
| Week 5 | 24 | 88.4<br>(± 110.3) | 9.0<br>(± 3.4) | 8.4M<br>(± 2.7M) | 10.0M<br>(± 3.0M) | P2 Solo |
| *Sum / mean* | 72 | 57.5<br>(± 48.7) | 7.4<br>(± 2.8) | 10.1M<br>(± 3.1M) | 9.6M<br>(± 2.8M) | |

**Table 3:** Mean cell-free DNA concentrations and sequencing reads obtained from the four evaluation cohorts.

| Evaluation cohorts | | Avg. cell-free DNA (± SD)<br>(ng/μl per animal) | | Avg. reads (± SD)<br>(million reads/animal) | | ONT device |
| --- | --- | --- | --- | --- | --- | --- |
|  | *n* | Control | Infected | Control | Infected |  |
| Pilot study | 22 | 28.2<br>(± 19.2) | 19.0<br>(± 15.5) | 1.6M<br>(± 0.26M) | 1.7M<br>(± 0.8M) | MinION |
| Specificity cohort -<br>*E. coli* | 12 | 4.0<br>(± 1.5) | 7.2<br>(± 3.1) | 4.7M<br>(± 1.3M) | 6.5M<br>(± 1.3M) | P2 Solo |
| Specificity cohort -<br>*G. anatis* | 12 | 9.0<br>(± 4.6) | 13.0<br>(± 2.2) | 5.0M<br>(± 1.9M) | 6.0M<br>(± 0.4M) | P2 Solo |
| Clinical & real-world<br>samples | 6 | † *gel* | 3.8<br>(± 2.4) | 2.0M<br>(± 1.4M) | 1.5M<br>(± 1.3M) | MinION |
| *Sum / mean* | 52 | 13.7<br>(± 8.4) | 10.7<br>(± 5.8) | 3.4M<br>(± 1.2M) | 3.9M<br>(± 0.9M) | |

† = samples were analysed with 2% agarose gel.

### Quality control of methylation sequencing of cell-free DNA regions

Median ONT sequencing coverages (± IQR) of cfDNA regions were 44.0x (± 40x), 214.0x (± 247x), 132.5x (± 87x), 49.0x (± 45) and 110.0x (± 90x) for the pilot study, long-term study, the specificity cohorts, the clinical samples and the real-world (zoo) samples, respectively (Fig. 4A). Full methylation sequencing statistics are presented in Supplementary file 6. Statistical comparison revealed that 14 out of 18 region classes throughout week 2, 4, and 5 exhibited significantly different methylation densities when compared between control vs infected (Fig. 4B).

**Fig. 4:**
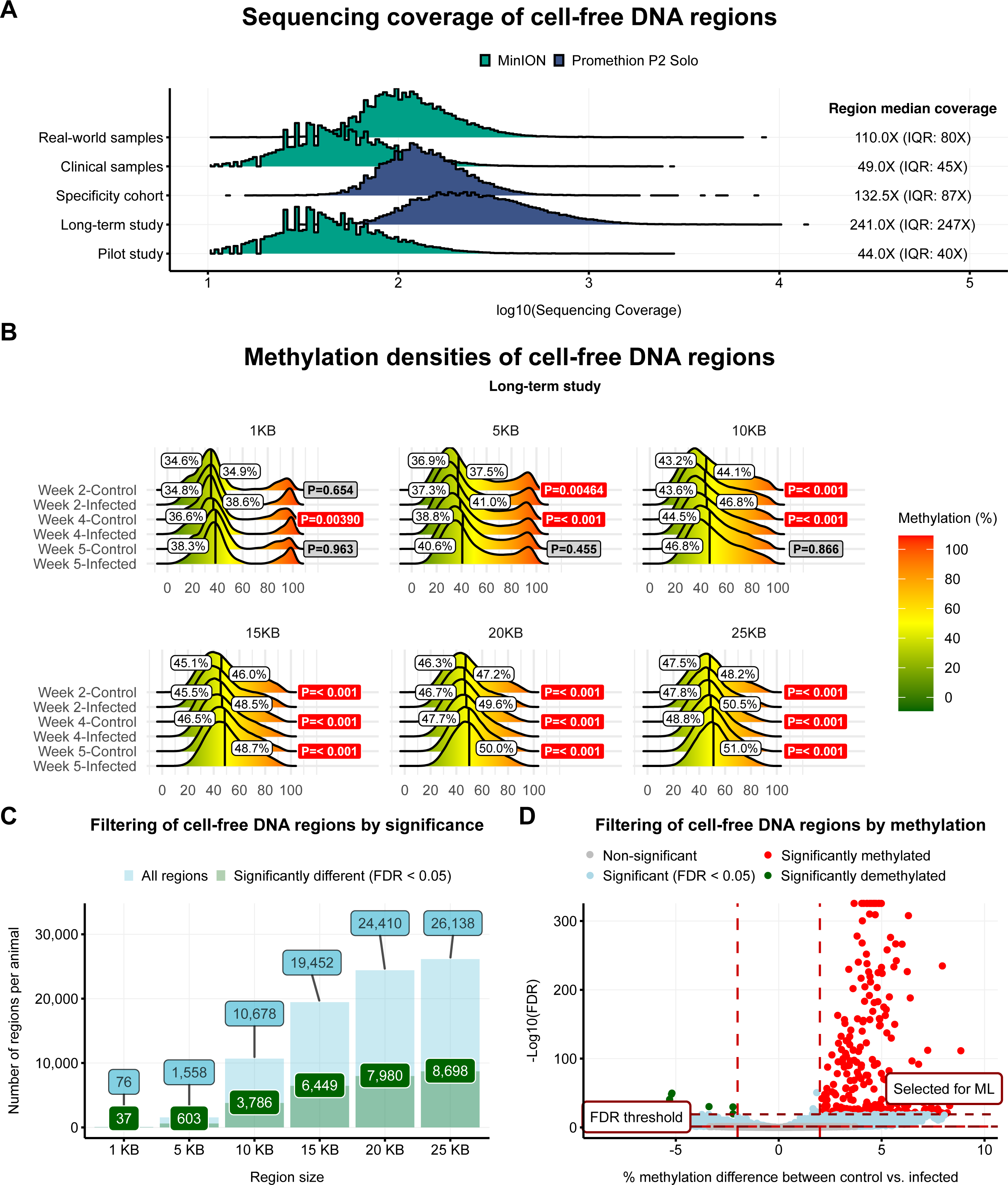
A) Log10 transformed sequencing coverage; B) methylation densities of the long-term study. White labels indicate median methylation. Red labels indicate statistical comparison between control vs infected groups; C) Filtering by *Af-*specific differentially methylated regions (DMRs) (green); D) Filtering by >2% methylation or demethylation. IQR: interquartile range; KB: kilobases.

### Filtering for *Af*-specific differentially methylated cell-free DNA regions

The number of DMRs ranged from 37 to 8,698 for the six region classes, respectively (Fig. 4C). Subsequently, these DMRs were subjected to a second filtering step, where only DMRs exhibiting a minimum methylation fold-change between control vs infected of either 2% or -2% methylation difference were selected for analysis (Fig. 4D). After methylation filtering, a total of 26 (1KB), 397 (5KB), 2986 (10KB), 5457 (15KB), 6782 (20KB) and 7280 (25KB) DMRs were saved and used as input markers for training ML models.

### Differentially methylated regions as marker sets and model training

Using a three-step filtering process, a total of 49 ML models passed the filters out of all 9,205 ML models and were assigned as candidate models whereas the remaining models were discarded. Three models were selected as the final diagnostic tests and named as follows: ‘High Accuracy test’ comprising 83 markers of 10KB each in a glmnet model, ‘Fast test’ comprising 22 markers of 10KB with an rf model, and ‘In situ test’ comprising 4 markers of 5KB with an lda model (Fig. 5A). Test statistics and marker genomic coordinates are found in Supplementary files 7 and 8. Methylation for each chicken across the markers of each diagnostic test, as well the three diagnostic tests themselves in the R package “caret” model format can found under data availability.

**Fig. 5:**
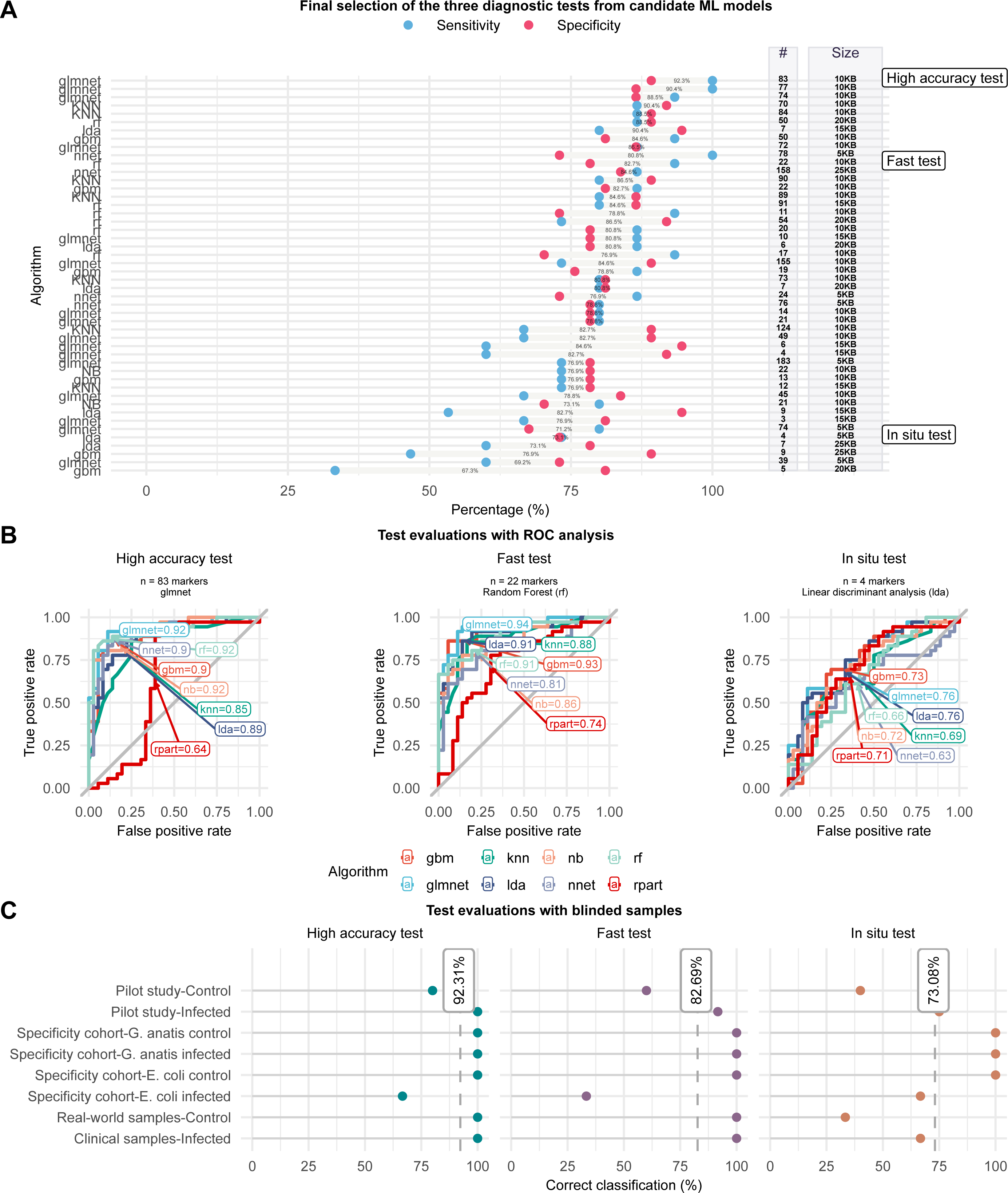
A) Final diagnostic test selection. Left Y-axis indicates ML algorithm, right Y-axes indicate number of markers (#) and their region class size; B) Area under curve (AUC) calculated from receiver operating characteristics (ROC); C) Correct classifications of samples from the performance cohort to their respective status (either *Af*-positive or *Af*-negative).

### Evaluation and final selection of the three diagnostic tests from candidate ML models

Test evaluation showed the area under curve (AUC) from receiver-operating characteristic (ROC) analysis to be 0.92, 0.91, and 0.76 for the High Accuracy, Fast, and In situ tests, respectively (Table 4). The algorithms for each of the models were glmnet, rf, and lda, respectively (Fig. 5B). The diagnostic accuracies were 92.3%, 82.7%, and 73.1%, corresponding to correct classifications of 48, 43, and 38 out of 52 blinded performance samples for the High Accuracy, fast, and In situ tests, respectively (Fig. 5C and Table 5). Specificity was lowest for *E. coli* experimentally infected animals, ranging from 33.4% to 66.7% correct classifications in the three tests, respectively. Specificities for control samples from the pilot study and the naturally infected clinical samples ranged between 40%-80% and 33.4%-100% for the three tests, respectively (Table 5).

**Table 4:**
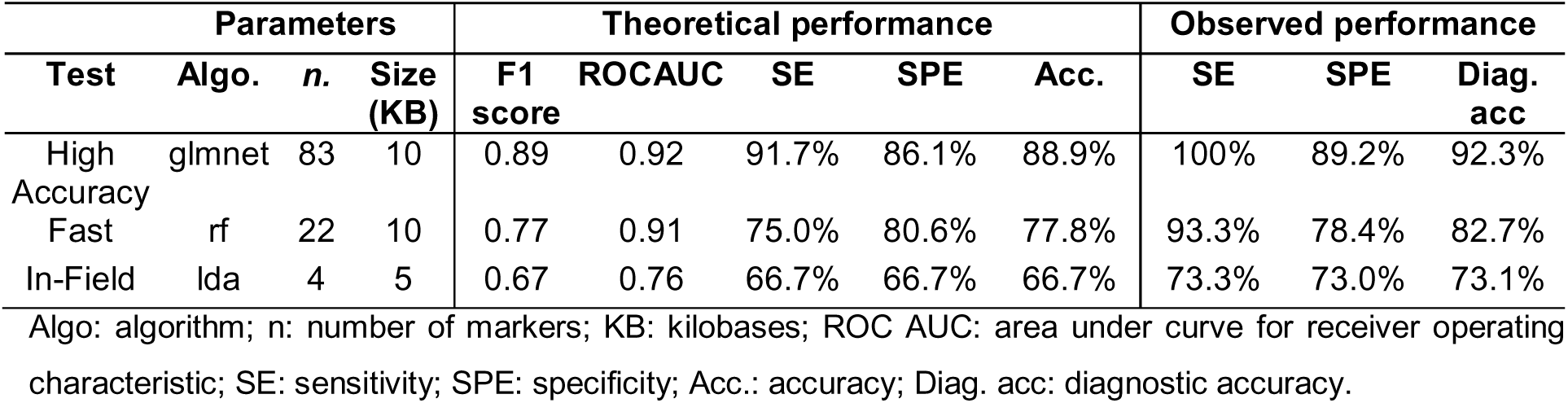
Performance of three final diagnostic tests obtained from theoretical evaluation of training samples and observed performance obtained from blinded evaluation samples.

**Table 5:** Final correct classifications of the three tests in details with sample sizes indicated for each cohort.

| Cohort |  | n | High Accuracy Test | Fast Test | In situ test | Mean % |
| --- | --- | --- | --- | --- | --- | --- |
| Correct classifications |  |  |  |  |  |  |
| Pilot study | Control | 10 | **80%** | 60% | 40% | 60% |
|  | Infected | 12 | **100%** | 91.7% | 75% | 89.2% |
| Specificity cohort – *E. coli* | Control | 6 | **100%** | 100% | 100% | 100% |
|  | Infected | 6 | **66.7%** | 33.3% | 66.7% | 55% |
| Specificity cohort – *G. anatis* | Control | 6 | **100%** | 100% | 100% | 100% |
|  | Infected | 6 | **100%** | 100% | 100% | 100% |
| Clinical & real-world samples | Control | 3 | **100%** | 100% | 33.3% | 76.7% |
|  | Infected | 3 | **100%** | 100% | 66.7% | 90% |
| Total correct: |  | 52 | **92.3%** | 82.7% | 73.1% | 82.7% |

n: number of samples.

### Individual diagnostic markers, functional annotations and reference probabilities

Median methylation in the High Accuracy test increased significantly (*p* < 0.05) in the long-term study week 5, the pilot study, the specificity cohort with *G. anatis*, and the real-world & clinical samples (Fig. 6A, Supplementary file 9). Functional annotation analysis showed overlaps with Epigenetically Modified Accessible Regions (EMARs), gene, transcript, enhancer, coding sequence (CDS), 5’ and 3’ untranslated regions (5/3’UTR), and exon regions. The four markers of the In situ test overlapped six features, the most overlapping features being EMAR, gene, transcript and enhancer regions (Fig. 6B).

**Fig. 6:**
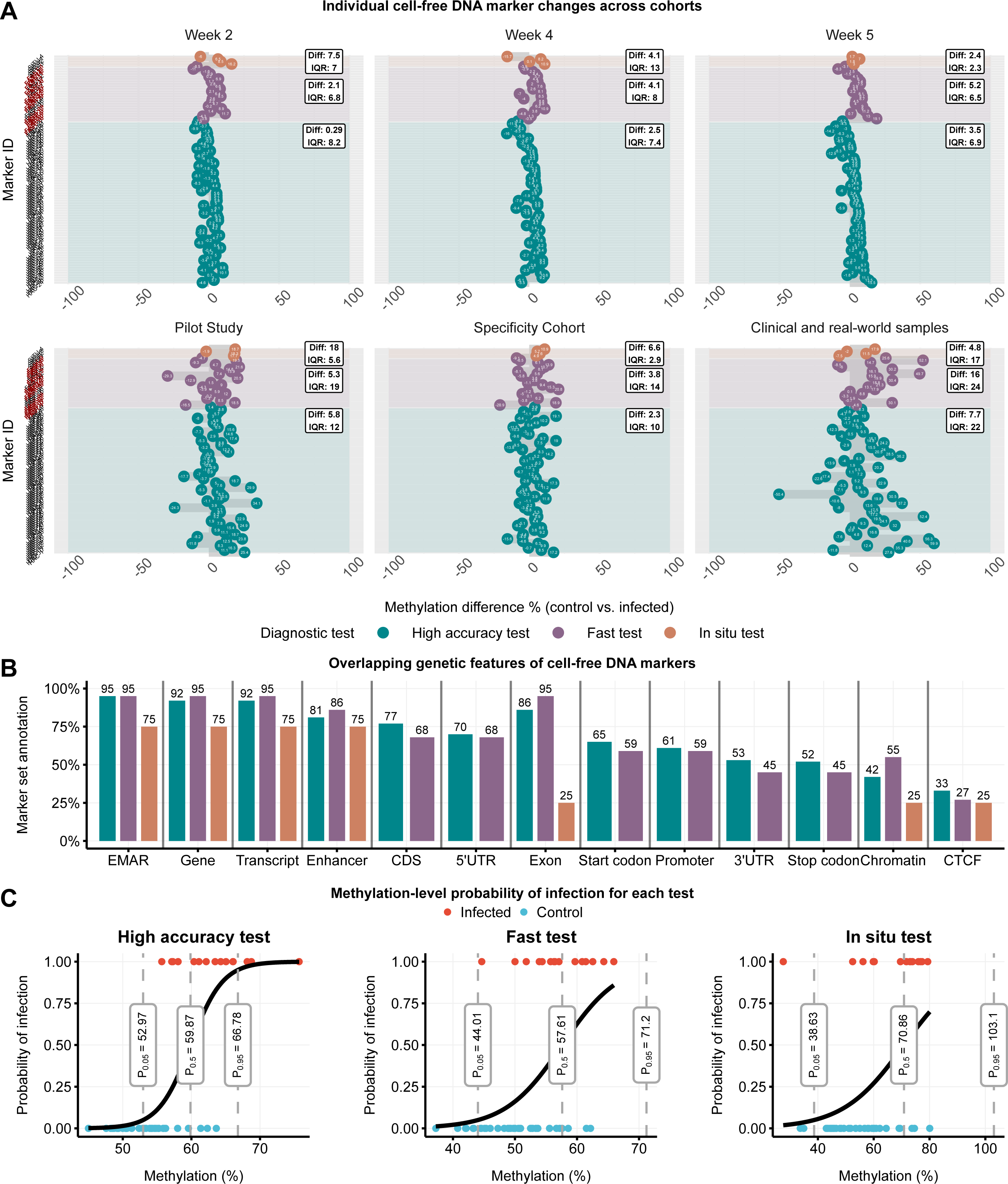
A) Marker methylation difference between control and infected. Red indicates duplicate markers; B) Marker overlaps of functional regions in the chicken genome; C) Methylation at 5% (P_0.05_), 50% (P_0.5_), and 95% (P_0.95_) probability of infection. UTR: Untranslated region; CDS: Coding Sequence, CTCF: zinc finger element; EMAR: Epigenetically Modified Accessible Regions.

The median methylation reference values were found to range between 49.1% to 70.7% in the High Accuracy test, 36.4% to 78.8% for the Fast test, and 20.6% to 121.2% for the In situ test (Table 6). For each test, the median methylation for a probability of P_0.5_, where the probability of a sample being either control or infection is equal, were found to be 59.9%, 57.6%, and 70.9% for the High Accuracy test, Fast test, and In situ test, respectively (Fig. 6C).

**Table 6:**
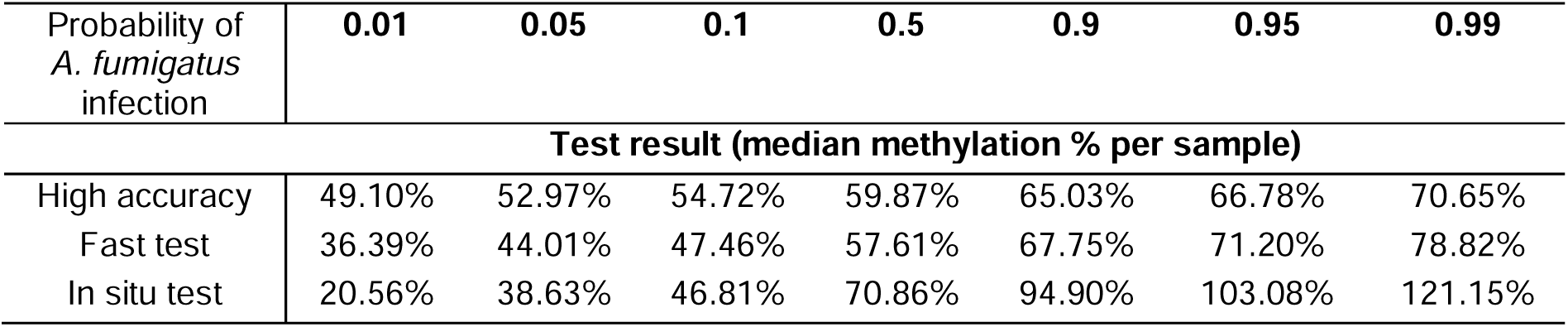
Probabilities of infection and their associated median methylation percentages for diagnostic test markers.

## Discussion

To the best of our knowledge, this is the first antemortem diagnostic test against avian *Af* infection based on the inference of cell death through infection-mediated changes in host cfDNA epigenetics. Using this strategy, we demonstrated a performance of up to 48 out of 52 (92.3%) correct classifications (SE: 100%, SPE: 89.2%) of blinded samples. The ML models were found to be highly robust as observed by ROC-AUC values of 0.92, 0.91 and 0.76, which are interpreted as excellent and fair by ROC assessment ^46,47^.

A major advantage of the cfDNA tests is the early detection of *Af* infection before clinical symptoms. In case of the infected pilot study samples, the High Accuracy test was able to correctly classify 100%, whereas the Fast test and In situ test classified 91.6% and 75% correctly. In contrast, other antemortem diagnostics were inconclusive and only SPE was able to discern significantly lower A:G ratios for the 50M group in the pilot study as well as the specificity cohort after infection. However, SPE is a non-specific technique and was also ambiguous; A:G ratios were significantly increased in the 5M group of the long-term study, and only the lowest dose (50K) was significantly decreased after the first week in the pilot study. Moreover, the inter-assay variability was good for TP (CV: 7.3%) but moderate for A:G ratios (CV: 17.9%), which for the latter is slightly above the generally accepted threshold for inter-assay CV at 15% ^48^, which could explain the variations observed. Antibody testing using IgG against *Af* did not identify any positive samples outside of the positive penguin control, regardless of time after exposure infection, thus exhibited moderate accuracy (64.6%). It might be explained by the chicks being day-old at the time of infection, as previous research showed critical age-related changes to the immune system the first 10 days post-hatching, where the primary immune response change from Th1 to Th17 ^49^.

Histopathological assessments showed that all tested infected samples from both *in vivo* studies were affected by one or more type of tissue pathology, whereas control samples were absent of pathology. However, not all samples were affected by all parameters; all 5M and 50M samples in the pilot study had oedema and presence of mononuclear cells, whereas the 50M also exhibited hyphae, granulomas and giant cells. The 5M samples in both pilot- and long-term study exhibited high BALT, but diagnosis of *A. fumigatus* infection would entirely depend on the emphasis placed on the individual parameters, and the comparative scoring system used. For the 5M groups, the presence of pathology ranged from 36.7%-55% for the pilot- and long-term study, respectively, compared to 85% (± 25.8%) for the 50M samples. These changes clearly highlighted the impact of increasing the dose concentration 10-fold and the seriousness of infection.

We hypothesise that the high sensitivity was derived from the perturbations in cfDNA methylation from the moment of increase in infection-mediated cell death, causing altered cfDNA shedding from infected tissues to the bloodstream ^50,51^ as well as shedding from dead immune cells ^32,52^. For the cfDNA pool in circulation, this led to an increase in subtle pathogen-specific epigenetic fingerprints, which we could resolve effectively due to the high sequencing coverage of DMRs. In a clinical scenario, adaptive sampling using a standard MinION flowcell ^53^ could analyse thousands of samples for the High Accuracy test, which requires a minimum yield of 8.3 million bases (MB) per sample. In practice, flowcells would be limited to 96 samples due to number of unique barcodes but still highly cost-effective. As NGS is becoming vastly more prevalent in clinical diagnostics and routine studies within animal- and veterinary sciences such as hereditary traits ^54^ and epigenetic programming ^55^, ONT sequencing could be a highly attractive choice for clinics and smaller labs without the need for additional specialised staff ^56^. Another option is to avoid NGS altogether using the In situ test where the four markers could be adapted to methylation-specific PCR ^57^.

Another major advantage of the technique proposed here analysing host cfDNA instead of microbial cfDNA (mcfDNA) by metagenomic NGS (mNGS) is the freedom of concern for environmental contamination. Around 90%-99% of cfDNA is from the host itself, whereas mcfDNA represent the remaining fraction, comprising 0.08%-4.85% bacteria, 0.00%-0.01% fungi, 0.00%-0.16% virus/phage origin ^58,59^. In human, a study by Kowarsky, et al. ^60^ conducted sequencing of 37 billion molecules from 1,351 samples of cfDNA, which revealed that an average of 0.45% cfDNA reads were of nonhuman origin, consistent with previous findings ^61^. Of these nonhuman reads, only approximately 1% could be identified in a curated microbiome database of almost 8,000 species of known pathogens ^60^, which underlines the sheer analytical difficulty in working with mcfDNA, especially for detecting early infection. This is further complicated by the probability of background contamination, as *Af* is an ubiquitous fungal mould in the environment ^62^, which can severely influence test results when using mNGS ^63,64^.

Previous research reported performance for detecting *Aspergillus* using mNGS or PCR on mcfDNA between SE: 53.0%-91.7% and SPE: 71.4%-97.0% ^65–70^. Thus, the High Accuracy test in the current study has a similar or even better performance in some cases (SE: 100% & SPE: 89.2%). This is a remarkable as the test is based on detecting the consequences of infection rather than detection of the pathogen DNA itself. We hypothesise that the shift in cell death dynamics during infection with either *Af, G. anatis* or *E. coli* resulted in methylation changes, which the ML models could resolve back to the causative pathogen in most cases. For the specificity cohorts, we observed that 100% of the *G. anatis* infected samples were classified correctly as non-*Af* infected, whereas the same number was 66.7% of the *E. coli* infected samples for the High Accuracy test. As none of the tests could correctly classify all infected *E. coli* samples, this leads to speculation that this infection shares similar cell death dynamics with *Af* infection. In this case, deeper (>44X) ONT sequencing coverage is needed to resolve the *E. coli* samples correctly.

ML-based analysis of cfDNA has already had its debut within the realm of cancer diagnostics ^50,71–73^. With regards to infectious diseases, a recent study found that active pulmonary tuberculosis (TB) led to higher cfDNA methylation levels, which was positively correlated with blood C-reactive protein (CRP) ^44^ in humans. Our findings agreed and were consistent with the observations that both global methylation of cfDNA and the test methylation markers of infected samples were found to be generally higher (Supplementary files 6 and 9) and positively correlated with the probability of *Af* infection (Table 6). Finally, the markers of the three tests overlapped to a large degree with EMARs (75%-95%), genes (75%-95%) and enhancer regions (86%-75%), emphasising possible functional roles and confirming the accuracy of the ONT guppy basecalling software for base modifications.

Limitations of this study include the sample size for real-world and clinical samples, which were limited, making it difficult to assess if any breed or age bias would affect the final diagnostic outcome. No bias was apparent in this study, but further studies should reaffirm that this is indeed the case. Moreover, it would have been interesting to measure acute phase proteins (APPs) like serum amyloid A (SAA) and α1-acid glycoprotein (AGP) ^74^, their dynamics and association with cfDNA and the innate immune response. Previous research found that *Af* can induce an acute inflammatory response through stimulation of SAA ^11^ and AGP plays a modulatory role in the immune response and appears to have a significant role in early stages of infection ^74^. Finally, larger trials should be conducted to assess and optimise the practical use and workflow of ONT sequencing for clinical diagnostics with an optimised pipeline for efficient diagnostic interpretation. In this context, specific APPs should be measured to provide greater sensitivity for changes in the acute phase response compared to SPE and to correlate with methylation dynamics of infection and cfDNA.

## Conclusion

The High Accuracy test for *Af* infection based on host cfDNA methylation markers demonstrated high performance (SE: 100%, SPE: 89.2%), diagnostic accuracy (92.3%), and robustness (ROC-AUC: 0.92), performing equally or better than current tests detecting *Af* using mNGS and PCR on mcfDNA, but with complete freedom of concern of false positives due to environmental contamination. Currently, the diagnostic tests are developed for use in poultry but has major potential if adapted to endangered avian species. This could be a breakthrough in the health monitoring of vulnerable species for conservation efforts.

## Materials and methods

### Animal model for *in vivo* studies

A full overview of the study design can be found in Fig. 1. Briefly, two overall cohorts were created: i) a model training cohort comprising a long-term *in vivo* study infected with *Af* (n = 60 animals), where a total of 72 cfDNA samples from three blood samplings were sequenced and used for ML modelling, and ii) a performance cohort comprising a pilot *in vivo* study (n = 63 animals), a specificity *in vivo* study (n = 24 animals), and clinical & real-world samples (n = 6 animals), where a total of 52 cfDNA samples were blinded and used to evaluate the ML models and the final diagnostic tests (Fig. 1).

Practically, all experimental animals (n = 123 individuals) were commercial day-old hybrid broiler chickens (Ross-308) procured from DanHatch A/S and housed at the Department of Veterinary and Animal Sciences, in a Biosafety Level 2 facility. (Frederiksberg, Copenhagen, Denmark). Upon arrival, the chicks acclimatised for 1 day before initiation of the study and throughout the two models, all chickens were provided with ad libitum broiler feed, fresh water, sand, and adequate heating. The temperature was monitored and kept consistently at 25°C (± 0.5°C) and daily care was provided by animal caretakers. All animal experiments were approved by the National Danish Animal Experiments Inspectorate (Dyreforsøgstilsynet), licence number (2019-15-0201-01611) and the Copenhagen Zoo Animal Care and Use Committee (1 October 2022). All animal experiments detailed in this study comply with the Animal Research: Reporting of In Vivo Experiments (ARRIVE) guidelines and EU Directive 2010/63/EU for animal experiments.

### Mycological cultures and preparation of inoculum

Stock culture of *Af* strain CBS112.33 was kindly provided by Centraalbureau voor Schimmelculture (CBS) (Utrecht, NL). Conidia were obtained and prepared by protocol from Thierry, et al. ^75^ with slight modifications. Stock culture was plated under sterile conditions on Sabouraud dextrose agar (SDA) plates and incubated at 25°C for 10 days. After incubation, the plates were transferred to sterile fume hood for conidia isolation. Conidia from each plate were harvested by sequential flushing with a total of 10 ml sterile PBS containing 0.01% (vol/vol) Tween-20, which was collected into 15 ml Falcon tubes and transferred directly to microscopy facilities. Subsequently, 10 µl gently inverted suspension from each tube was carefully counted in a Fuchs-Rosenthal counting chamber by light microscopy at 400x magnification.

Tubes with the enumerated conidia were pelleted by centrifugation at 4000 x *g* for 30 min at 4°C, supernatant discarded, and finally, the volumes were collected and resuspended into a master suspension of 5 x 10^8^ conidia in 10 ml sterile PBS. The master suspension was used to create a dilution series of with dose concentrations of 5 x 10^7^, 5 x 10^6^, 5 x 10^5^, and 5 x 10^4^ conidia in a 100 µl volume, from now on designated 50M, 5M, 0.5M, and 50K. These four final suspensions were stored at 4°C and used as inoculum for experimental animal inoculations. To ensure the quality of the conidia, the whole procedure was always conducted the day before the start of a new animal experiment.

### Quality control of infection doses by semi-quantitative real-time PCR

To determine the consistency and accuracy of the two inocula created for the pilot study and the long-term study, as well as ensuring the dose concentrations, a TaqMan probe-based real-time quantitative PCR (qPCR) assay was used in conjunction with plasmid DNA to accurately assess the conidial equivalents ^76^ following the procedure by Melloul, et al. ^77^. This procedure takes advantage that the *FKS* gene is a single-gene in the *Af* genome. As conidia are haploid units ^78^ and thus only carries a single *FKS* copy, the number of conidia can be determined by semi-quantitative standard curve using plasmid DNA with known molarities. Briefly, 100 µl of the prepared inocula corresponding to an individual chicken dose was used as input material for the Qiagen DNeasy Blood & Tissue kit (Qiagen, Hilden, GmbH) following manufactures instructions for a total elution volume of 200 µl. Genomic DNA (gDNA) was quantified using a NanoDrop One UV/Vis spectrophotometer (ThermoFischer, Denmark). Conidial equivalents were quantified by primers and probes for the single-copy *FKS* gene, previously reported by Costa, et al. ^79^: Sense amplification primer, AFKS1 (5′-GCCTGGTAGTGAAGCTGAGCGT-3′), antisense amplification primer, AFKS2 (5′-CGGTGA ATGTAGGCATGTTGTCC-3′) and the TaqMan *FKS* probe (5′-6-FAM-TCACTCTCTACCCCCATGCCCGAGCC-BHQ1-3′). For semi-quantification by standard curve, we constructed a 490 bp long synthetic DNA fragment with known molarity of the *FKS* gene (GenBank accession: U79728, from bp 2731 to bp 3211, Supplementary File 1), which was procured from Invitrogen GeneArt Synthesis (ThermoFischer, Denmark) and diluted into five log^10^ concentrations. The final reaction mixtures were: 2 µl DNA, 0.9 µM sense and antisense primers, 0.2 µM FKS TaqMan probe, 10 µl TaqMan™ Fast Advanced Master Mix for qPCR (Applied Biosystems™, Roskilde, Denmark) and 4 µl nuclease-free H_2_O in 20 µl reaction volume. The qPCR reactions were run using a LightCycler® 96 detection system (Roche, Meylan, France) under the following conditions: 50°C for 2 min, 95°C for 10 min, then 50 cycles of 15 s at 95°C and 1 min at 65°C ^77^. All data was analysed with LightCycler^®^ 96 Application Software (v. 1.1.0.1320, Roche, Switzerland) and exported to Microsoft Excel. Amplification efficiency was calculated as e = 10^−1/slope^. All qPCR experiments were reported in accordance with the MIQE guidelines ^80^. Finally, median (± IQR) *FKS* gene copy counts were compared statistically with the enumerated microscopy counts and between similar dose concentrations between the two *in vivo* studies using a Kruskal-Wallis test.

### *In vivo* studies: Pilot study for *Af* dose determination

A total of 63 one-day-old chickens divided into five groups were used to determine correct concentrations of the *Af* inoculum. After one day of acclimatisation, each of the five groups was subjected to intra-tracheal infection of 100 µl PBS containing 50K, 0.5M, 5M or 50M conidia or only sterile PBS. Monitoring of the animals was done intensively, with hourly visual health and behavioural checks for the first three days with exception of 8 h nightly breaks. In accordance with the Danish Animal Experiments Inspectorate guidelines ^81^, a maximum of 1.5 ml blood was drawn for serum protein electrophoresis (SPE) and cfDNA sequencing from the brachial vein on day 8 (week 1), 15 (week 2), and 23 (week 3). Subsequently, all animals were weighed and assessed visually. Blood from four additional eggs served as extra negative controls before the pilot study trial started, of which blood was drawn directly through the shell one day before hatching prior to euthanasia by rapid cervical dislocation. Serum was obtained by letting blood samples coagulate overnight at room temperature and subsequent centrifugation at 2000 x G for 15 min at 4°C.

### *In vivo* studies: Long-term study for model training

A total of 60 one-day-old chicks were divided into two equally sized groups and housed in separate rooms in temperature- and humidity-controlled isolation facilities. After one day of acclimatisation, all chickens were dosed intra-tracheally with 100 µl sterile PBS containing either 5M conidia or only sterile PBS. Blood for SPE and cfDNA sequencing was drawn from the brachial vein on day 15 (weeks 2), day 29 (week 4), and day 35 (week 5) and all animals were weighed. Serum was obtained by letting blood samples coagulate overnight at room temperature and subsequent centrifugation at 2000 x G for 15 min at 4°C.

### Specificity, clinical samples

The specificity cohort comprised serum samples (n = 24) from two previously conducted longitudinal studies with *E. coli* infection (Bojesen et al. 2025, in prep) and *G. anatis* infection (Antenucci, et al. ^82^) in chickens. Briefly, the *E. coli* group comprised a total of 12 serum samples from six chickens (breed: Ross-308) infected intra-tracheally with 1 x 10^6^ live colony-forming units (CFU) per chicken of *E. coli* (strain DH269, sequence type (ST) 428) in a longitudinal study design: blood draws were conducted on January 23, 2024, immediately before intra-tracheal infection, and from the same individuals on January 25, 2024 or two days post-infection (dpi). The *G. anatis* group comprised a total of 12 serum samples from six chickens (breed: 16-weeks old Lohmann-Brown layer chickens) infected intra-peritoneally with 10^7^ live CFU per chicken of *G. anatis* (strain: 10672/9) in a longer longitudinal study design: blood draws were conducted on April 16, 2019, and again from the same individuals on May 6, 2019 or 21 days dpi (Antenucci, et al. ^82^).

Serum samples from clinical cases were shared kindly by Laboklin GmbH. diagnostic facility (Bad Kissingen, Germany), and were used as unrelated suspected positive samples. Prior to this study, each sample was tested using an Indirect Haemagglutination Assay (IHA) using the Hemkit Aspergillus IHA (Ravo Diagnostika GmbH, Germany). Serum was taken on 28 December 2023, 16 October 2023, and 30 September 2023, with IHA titres of 1:80, 1:160, and 1:320 respectively. Finally, three serum samples from clinically healthy adult chickens were obtained from the Copenhagen Zoo and were used as suspected negative blinded samples. Each of the three samples were taken on January 21, 2010, and serum had been stored at -80°C since collection.

### Serum protein electrophoresis – *in vivo* studies and specificity cohort

To assess the acute phase proteins (APPs), the humoral immune response, and general health condition in the *in vivo* chicken models and performance cohort, SPE was conducted to obtain albumin/globulin (A/G) ratios. Total protein in g/dL (TP) was measured in 2 µl serum for each sample using a standard veterinary refractometer (Atago T2-Ne Clinical, Atago Co., Ltd, Japan) at standard atmospheric pressure and room temperature. SPE was conducted for each serum sample by agarose gel electrophoresis (AGE) using the EP001 Kit (DDS Diagnostic S.R.L, Bucharest, Romania) following manufactures instructions. The gels were loaded and run in a standard Bio-Rad PowerPac Basic gel electrophoresis apparatus (Bio-Rad, CA, USA) at 100V for 20 min. After fix, stain, and bleach steps, the dried gel was scanned using an Epson V550 (Epson, USA) with SilverFast SE 8.8 (LaserSoft, Kiel, Germany). An electropherogram was generated for each sample using the gel analysis function in ImageJ software (v. 1.54d, NIH, USA ^83^). Finally, the area under curve was computed for each of the fractions (prealbumin, albumin, α1, α2, β, and γ) and A:G ratios were calculated by division of the summed prealbumin and albumin fractions by the sum of α-, β-, and γ-globulins as per standard convention ^84^. To test and assess the inter-assay variation, 10 samples from five animals were analysed twice, each at different dates. Subsequently, the CV was calculated for TP and A/G ratios by conventional SD divided by the mean.

### Postmortem gross examination

Euthanasia was conducted by rapid cervical dislocation. To assess pathological changes in the respiratory tissues and organs, necropsies of each individual bird was conducted within 1 h of euthanasia. All animals from the pilot study (n = 63) were subjected to postmortem examinations, comprising the groups 50K (n = 13), 0.5M (n = 13), 5M (n = 15), 50M (n = 10) and the control group (n = 12). For the long-term study, all animals (n = 60) were subjected to postmortem examinations, comprising the groups 5M (n = 30) and control group (n = 30). A pre-developed necropsy scoring sheet was used to evaluate the peritoneum, left airsac, left lung (with weight), spleen (with weight), liver, and trachea. Each organ was evaluated on parameters such as inflammatory reaction, amount and type of exudate, as well as transparency of the airsac, congestion, proliferation of the spleen, granuloma formation, and other anomalies. Scores ranged from 0 to 4, where 0 represented absence of lesions and 4 was marked lesions. Scoring definitions for each parameter is available in Supplementary File 10.

To compare scorings statistically, scores for each parameter (Inflammatory reaction, exudate etc.) in each tissue (Airsac, lung etc) was summed by each dose group (50K, 0.5K, 5M, 50M, control) and divided by the number of animals in the dose group to obtain weighted scores. By summing all weighted scores from all control samples in both *in vivo* studies and dividing by the total number of control animals (n = 42) multiplied by the total amount of tested parameters (n = 15), a baseline control frequency was obtained. Using a binomial test, this frequency was used to compare summed scorings from individual parameters for each infected dose group as the number of successes with the total number of animals examined for each parameter in each tissue as the number of trials. Finally, the p-values were subjected to multiple testing correction using Benjamini-Hochberg (FDR) correction.

### Histopathology and immunohistochemistry on lung tissue

After each necropsy and scoring, one piece of the left lung was fixed in formalin for histology. Briefly, following paraffin-embedment tissue sections of 4 µm in thickness were stained with haematoxylin and eosin (HE). Immunohistochemistry (IHC) was conducted by mounting on positively charged glass slides (Superfrost^®^plus, Thermo Scientific, Germany) and subsequently, the slides were subjected to deparaffinisation, rehydration and antigen retrieval (95°C for 30 min). The immunostaining was conducted with a primary antibody against *Af* (Anti-Aspergillus Antibody [WF-AF-1] (A281971)) and detected using UltraVision LP Detection System HRP according to the manufacturer’s instructions. The fungal elements of *Af w*ere stained red. Finally, histopathological evaluations were conducted for 12 parameters for a total of 15 samples from the 50M (n = 5) and 5M (n = 10) groups in the pilot study, and a total of 20 samples from the 5M (n = 10) and control (n = 10) groups from the long-term study. Evaluations were compared statistically with non-parametric Wilcoxon tests with Benjamini-Hochberg multiple testing correction (FDR) and visualised using the ggstats (v. 0.7.0) ^85^ R package.

### Serology-based diagnostics using chicken IgG

To assess *Aspergillus* specific antibody levels, the LDBio Aspergillus IgG kit (LDBio, Lyon, France) western blot kit was used in conjunction with a chicken-IgG conjugate (LDBio, France). All determinations were done according to manufactures instructions. Positive controls comprised serum obtained from an *Af* positive penguin confirmed by necropsy, which was collected in January 2023 in the Copenhagen Zoo before euthanasia. The final blots were scanned with an Epson v550 (Epson, USA) using SilverFast SE 8.8 software for confirmation of minimum two bands for a positive test in accordance with manufactures instructions.

### Final definitions of the training and the performance cohort

In total, 124 samples were selected for cfDNA ONT sequencing. For the training cohort, a total of 72 samples from the long-term study were selected, comprising samples from week 2 (n = 24), week 4 (n = 24), and week 5 (n = 24), equally divided between infected (5M) and control groups. For the performance cohort, a total of 52 samples unrelated to the long-term study was designated as unknown blinded samples for evaluating the diagnostic performance of each ML model. The unknown blinded samples comprised of the pilot study for dose determination (n = 22), which were infected samples (n = 11 from 5M and n = 1 from 50M groups) and healthy controls (n = 10), the *G. anatis* and *E. coli* specificity samples (n = 24), three clinical samples (n = 3) and three real-world archived samples from the Copenhagen Zoo (n = 3). A total of 52 samples were used as performance cohort and evaluation.

### Cell-free DNA extractions and quality control

A total of 124 serum samples from the training (n = 72) and performance cohorts (n = 52) were selected for cfDNA extractions and ONT methylation sequencing. The training cohort comprised samples from the long-term study (n = 72), and the performance cohort comprised samples from the pilot study (n = 22), two specificity studies (n = 24), and clinical & real-world samples (n = 6). A total of 200 µl serum per sample was used as input for cfDNA extraction with the Norgen Biotek Circulating Cell-Free DNA Micro kit (cat. no. 55500) following manufacturer’s instructions. Before extractions, all serum samples were subjected to a centrifugation step for 2 min at 400 x g (∼2,000 RPM) prior to extraction. After cfDNA extraction, all samples were measured on a Bioanalyzer 2100 (Agilent, Germany) using the High Sensitivity DNA system (cat. no. 5067-4626) following manufacturer’s instructions. All cfDNA samples were stored at -80°C. No cfDNA concentrations were available from clinical & real-world control group as these samples were checked by 2% agarose gel.

### ONT sequencing of cell-free DNA from training and performance cohorts

All ONT sample preparation and flow cells were performed using the newest version 14 chemistry. The DNA libraries for sequencing were prepared using the Native Barcoding Ligation Sequencing kit (SQK-NBD114.24) with a sample input volume of 11 µl cfDNA including 1 µl diluted control DNA (DCS). The sequencing protocol used was the Native Barcoding gDNA 24 (v14-sqk-nbd114-24-NBE_9169_v114_revQ_15Sep2022). All steps were done according to manufactures instructions, except for the modification that all bead clean-up steps were multiplied by a factor of 2.5x as recommended by Martignano, et al. ^86^. All training samples, including the specificity cohort, was sequenced with the Nanopore P2 Solo sequencer. The pilot study as well as clinical and in-house samples were sequenced with the ONT MinION Sequencer. All flow cells, both PromethION and MinION flow cells, were of type R10.4.1 and each individual cell was checked before runs for adequate pore numbers. Raw sequencing data in pod5 format was transferred directly to a Samsung T7 external SSD after each run.

### Basecalling, cell-free DNA read mapping, and methylation data

All basecalling, mapping, and extraction of base modification (methylation) data was done on the national life-science high performance computer cluster (HPC) Computerome (DTU, Risø, DK) and the Aarhus University Genome DK Cluster (Aarhus, DK). First, ONT reads were transferred to the HPC and raw pod5 files were basecalled and respective cytosine base modifications (5mC and 5hmC) were computed using the ONT guppy (v. 6.5.7) software in High Accuracy mode using setting the barcode_kits flag to ‘SQK-NBD114-24**’** and using the model file ‘dna_r10.4.1_e8.2_400bps_modbases_5mc_cg_hac.cfg’. Basecalling was accelerated with an Nvidia Tesla V100 GPU. Subsequently, quality assessment of sequencing runs were evaluated using MinIONQC (v. 1.4.2) ^87^. After quality assessment, all cfDNA reads together with base modifications were compiled together in an unaligned BAM file for each sample (or chicken). Each unaligned BAM file was mapped to the Ensembl *Gallus gallus* reference genome (bGalGal1.mat.broiler.GRCg7b rel. 108) ^88^ using samtools fastq converter (v. 1.16) ^89^ piped together with minimap2 (v. 2.24-r1122) ^90^ in ONT mode. Mapped reads were sorted and indexed using samtools and evaluated using qualimap (v. 2.2.1) ^91^. Finally, the aligned BAM files with base modifications (methylation patterns) were used as input for the ONT modkit (v. 0.1.2) pileup software in “cpg combine mods”, which extracted all relevant base modifications (methylation patterns) for each sample into a bed file, which were used for further analysis. The full bash script pipeline for raw Nanopore pod5 basecalling, reference genome mapping, methylation extraction, and bed file generation is available in Supplementary file 11.

### Statistical processing, filtering, and cfDNA region size generation

For statistical processing, each bed file was downloaded and used as samples for the R package methylKit (v. 1.24.0) used with R (v. 4.4.1) ^92^ and Rstudio (v. 2024.04.1+748) ^93^. When loaded into methylKit using methRead, all methylation bed files were checked for irregularities and finally, normalised for read coverage by their median CpG counts using the methylKit normalizeCoverage() function and the methylObject was ready for binning the cfDNA into six region size classes. Using the bedtools (v. 2.31.1) makeWindows software, the Ensembl chicken reference genome was divided into sequential unbiased non-overlapping genomic regions, which started from chromosome 1 and ended in the last position of the sex and mitochondrial chromosomes. Six individual genome partitions were made, which comprised genome-wide regions of 1 KB, 5 KB, 10 KB, 15 KB, 20 KB, and 25 KB, and each genome partition was saved as an individual bed file. The normalised methylation percentages generated earlier in the methylObject were binned into their regional methylation counts using the genome partitions generated previously using the regionCounts() function in methylKit, which was repeated six individual times for each partition (1, 5, 10, 15, 20, and 25 KB). Herein, a filtering step was included, that required a minimum coverage of 10X cfDNA reads over each region for each sample for a region to be saved for further statistical analysis. This step reduced the number of regions from the millions to the thousands to conserve only highly covered filtered regions for appropriate statistical testing with highest power and sequencing accuracy.

### Filtering for *Af*-specific differentially methylated cell-free DNA regions

Differential methylation was conducted on the highly covered filtered regions using methylKit. Briefly, statistical comparison was done with a logistic regression test, where methylation proportions (the numbers of methylated cytosines and unmethylated cytosines for each region) was compared statistically throughout the two groups (5M infected vs controls). If the fraction of methylated cytosines were significantly different across infected and control groups, the region was defined as a *Af-*specific DMR. Full details of the procedure can be found in Akalin, et al. ^94^. Finally, Benjamini-Hochberg multiple testing correction was applied to all DMRs and regions, which were significantly (FDR < 0.05) different between infection and controls were saved as potential markers and used as input into subsequent training of ML models. The markers were plotted using ggpubR (v. 0.6.0) ^95^, ggplot2 (v. 3.4.3) ^96^, ggridges (v. 0.5.6) ^97^, ggrgl (v. 0.1.0) ^98^, patchwork (v. 1.3.0) ^99^ and Pheatmap (v. 1.0.12) ^100^ in R. Methylation regions from individual animals from the long-term study was selected on the two criteria of being significantly differently methylated or demethylated (FDR < 0.05) and demonstrating a numeric median methylation change of at least 2% or -2% when the region was compared between control and infected groups from samples of week 2, week 4, and week 5 of the long-term study. DMRs passing this two-stage filtering process were selected for input as markers for training the ML models.

### Model training with machine learning (ML) algorithms

After the two-stage filtering, the top significant DMRs and their associated methylation data in the training cohort (n = 72) were used as markers for training the ML models. Using the caret (v. 6.0-94) ^101^ package in R, eight commonly applied ML algorithms were trained with the train() function with different combinations of the DMRs described below and their associated methylation data extracted from the training cohort, creating matrices of 72 samples with methylation data for a variable number of markers. The performance of the trained ML model and its algorithm was evaluated by the methylation data from the blinded performance cohort (n = 52) using the predict() function. The eight algorithms tested were: K-nearest neighbour (knn), Lasso and elastic-net regularized generalized linear models (glmnet), gradient boost machine (gbm), Recursive Partitioning and Regression Trees (rpart), random forest (rf), linear discriminant analysis (lda), naïve Bayes (nb), and neural network (nnet). A training control object was set up using the function trainControl() in the caret package for controlling resampling. The method used was repeated cross-validation (“repeatedcv”) using a 10-fold cross validation, repeated 5 times (3 times for rf and nb), and set to output class probabilities (classProbs = T) and to save the predicted values (savePredictions = T).

To find the optimal number of markers to include in a model and their respective sizes as previously described, a permutation of test parameters was set up to shuffle the markers in search for ideal combinations. Overall, the permutation parameters to be satisfied were: 1) twelve marker sets comprising markers ranked by significance, with sets ranging from being 10 markers to 285 markers long in increments of 25 markers; 2) additional filtering by methylation/demethylation up to 10% (in steps of 1% increments), and 3) markers sizes between 1, 5, 10, 15, 20, and 25 KB. The parameters led to 6,803 different combinations of marker sets with different sizes and methylation differences. All combinations were used as input to train eight ML algorithms, which led the total and final unique number of trained ML models to be 9,205 models for testing and evaluation.

### Evaluation of the diagnostic tests with the blinded performance cohort

All 9,205 ML models were evaluated by their ability to correctly predict the infection status of the blinded performance cohort. To find the most promising models, a set of filtering criteria were established by analysis and careful curation of the prediction performance and models were saved by passing at least one criterion: i) higher than 90% accuracy, ii) higher than 85% accuracy with fewer than 50 markers, iii) or higher than 75% accuracy with fewer than 10 markers of maximum 5 KB sizes. Finally, filtered models were evaluated with MLeval R package (v. 0.3) ^102^ and models having an area under the curve (AUC) value of over 0.75 were saved for final evaluation.

### Final selection of the three diagnostic tests from candidate ML models

All ML models that passed the subsequent filtering steps using the performance cohort were evaluated by hand curation on parameters of theoretical and observed performance characteristics calculated by the R Package MLEval (v. 0.3) ^102^. Final selection of the models for the three tests were selected on a list of prioritised criteria: 1) diagnostic accuracy, 2) sensitivity, 3) specificity, 4) area under curve (AUC) from receiver operating characteristic curve (ROC) analysis, 5) number of markers, 6) region size class of the markers, and 7) number of specificity samples that were wrongly classified.

### Individual diagnostic markers and their functional annotations

Summary statistics, chromosome mapping, genomic locations, and functional annotation was conducted with the R packages BRGenomics (v. 1.12.0) ^103^ and genomation (v. 1.36.0) ^104^. Function annotation of features such as genes, transcripts, exons were conducted using the Ensembl chicken feature genome (bGalGal1.mat.broiler.GRCg7b release 108). Functional analysis was conducted by comparison of diagnostic markers with 13 known genetic, transcriptional, and epigenetic activity revealed that the High Accuracy and Fast test contained markers, which overlapped to some degree all features. For regulatory features and Epigenetically Modified Accessible Regions (EMARs), the Ensembl chicken regulatory features reference genome was used (bGalGal1.mat.broiler.GRCg7b release 113).

### Reference methylation values and infection probabilities for each diagnostic test

Reference values for each diagnostic test (High Accuracy test (n = 83 markers), Fast test (n = 22 markers), In situ test (n = 4 markers) was calculated by logistic regression using a binomial glm model in R and comparing the median methylation values per sample for each marker set with the true status of each sample. Using logistic regression, the reference methylation values and intervals for clinical interpretation of new samples without running the full ML models were calculated for each of the three diagnostic tests for probabilities of 0.01, 0.05, 0.10, 0.50, 0.90, 0.95 and 0.99 of having infection with *Af*.

## Acknowledgments

We are deeply grateful to Tuomas O. Kilpeläinen, Jonathan R. Belanich and Boglár Gál at the Novo Nordisk Foundation Center for Basic Metabolic Research (CBMR), University of Copenhagen (DK), and Tom Gilbert and Sarah Mak at the GLOBE Institute at University of Copenhagen (DK) for providing access to Oxford Nanopore Promethion sequencing facilities and expert assistance. We sincerely thank and acknowledge Natascha Emilie Schaleck for excellent Nanopore MinION sequencing, as well as Jens H. Madsen and Linett Rasmussen for laboratory assay validation and cell-free DNA sample quality control. Moreover, we are deeply grateful for highly skilled postmortem work done by Rebecca Kristiane Dalgaard Thacker, Marie Palmqvist, Katrine Wedel Aagaard, Rimsha Farooq and Qian Zhou, and to Simon Bruslund, Bernhard Drag and Sigrid Stokbro Hess for important feedback on avian biology, method optimisation and statistical analysis. We are deeply grateful to the skilled animal keepers and veterinarians at the Department of Veterinary and Animal Sciences at University of Copenhagen (DK) for skilled assistance in animal keeping and blood sampling, and the Conservation and Veterinary Team at the Copenhagen Zoo for skilled assistance and consultations regarding clinical assays. We kindly acknowledge Laboklin Diagnostic Facility (DE) for sharing samples and validation results with us, as well as the Division of Comparative Pathology, Department of Pathology & Laboratory Medicine, University of Miami Miller School of Medicine (USA) for assay consultation and Technical University of Denmark (DK) and Aarhus University (DK) for granting us access to the high-performance server clusters Computerome and GenomeDK. Finally. we acknowledge InnovationFund Denmark, Benzon Foundation, Loro Parque Foundation, Association of Avian Veterinarians (AAV), European Association of Avian Veterinarians (EAAV) as well as Familien Hede Nielsen Foundation and H.P. Olsen og Hustrus Mindefond for funding of this project.

## Author contributions

MHD prepared and conducted animal sampling and experiments, designed and conducted the laboratory, ONT sequencing, and machine learning diagnostic workflows and wrote the first draft of manuscript; CH prepared study design, validated the statistical analysis, provided project management and oversight, and provided scientific feedback; LLP provided veterinary assistance, animal sampling and consultancy for the animal experiments; HEJ conducted histological evaluations and slide photography; SAT provided veterinary consultancy in clinical microbiology and fungal culturing; CL and CC validated cell-free DNA and serum protein electrophoresis assays and provided assay development feedback; MFB and AMB designed the study, provided project management and oversight, as well as veterinary and microbiological consultancy. All authors have approved the final draft of the manuscript.

## Data availability

All diagnostic models for the **High Accuracy test**, the **Fast test**, and in the **In Situ** test are available in original caret format for R (.rds), as well as processed methylation files for each sample (n = 124) in bed format (without duplications) is available through Zenodo:

DOI: **10.5281/zenodo.15194046**

All ONT raw sequencing data in fast5 libraries supporting the findings are available through the European Bioinformatics Institute (EBI), European Nucleotide Archive (ENA):

Accession no.: **PRJEB86769**

https://www.ebi.ac.uk/ena/browser/view/PRJEB86769

For basecalling and demultiplexing the libraries, sample barcode lists can be found in Supplementary file 6.

## Animal ethics statement

All animal experiments were approved by the National Danish Animal Experiments Inspectorate (Dyreforsøgstilsynet), licence number (2019-15-0201-01611) and the Copenhagen Zoo Animal Care and Use Committee (1 October 2022). All animal experiments detailed in this study comply with the Animal Research: Reporting of In Vivo Experiments (ARRIVE) guidelines and EU Directive 2010/63/EU for animal experiments.

## Supplementary files

**Supplementary file 1:** Synthesised DNA fragment in FASTA format used for RT-qPCR *FKS* assay for quantification of molecular conidial equivalents (CE) and conidial equivalents (CE) obtained from the *FKS* gene assay compared to enumerated microscopy of *Aspergillus fumigatus* conidia for the *in vivo* studies.

**Supplementary file 2:** Serum protein electrophoresis (SPE) data used for calculating albulin:globulin (A:G) ratios for all samples tested from the *in vivo* studies, with all raw calculations included. Only animals with lab IDs were included in the cell-free DNA study.

**Supplementary file 3:** Full phenotype dataset including pathogen information for the training cohort comprising the long-term *in vivo* study with all postmortem scorings included.

**Supplementary file 4:** Full phenotype dataset including pathogen information for the performance cohort comprising the pilot *in vivo* study, specificity cohort, clinical- and real-world samples with all postmortem scorings included.

**Supplementary file 5:** All results from antibody testing using the LDBio IgG kit for all cohorts.

**Supplementary file 6:** Median methylation (percentages ± IQR) of the six size classes of regions for each study and cohort, as well as the numbers of regions included for analysis.

**Supplementary file 7:** Final filtering of machine-learning (ML) models as candidates for the diagnostic tests. The three final selected tests are shown in the top three rows.

**Supplementary file 8:** Differentially methylated cell-free DNA regions (DMR) as diagnostic markers included in each of the three tests.

**Supplementary file 9:** Median methylation values based on infection status with statistical comparison using Wilcoxon.

**Supplementary File 10:** Post-mortem scoring sheet with scoring grading definitions for each individual parameter.

**Supplementary File 11:** Bash script pipeline for basecalling, mapping/alignment, and methylation extraction in BED-file format of raw ONT Nanopore sequencing data.

