## Supplementary file 1 for "New high accuracy diagnostics for avian *Aspergillus fumigatus* infection using Nanopore methylation sequencing of host cell-free DNA and machine learning prediction"

Synthesised DNA fragment used for RT-qPCR *FKS* assay for quantification of molecular conidial equivalents (CE).

**GenBank accession ID:** U79728, from bp 2731 to bp 3211

**FASTA**:

AGGTTCTCGCCACAACCGACATGGAAATCAAGTACAAGCCTAAGGTATTGATTTCTCAAGTTTGGAATGCCATCATCATCTCAATGTACCGGGAGCATCTGCTGGCTATAGACCATGTTCAGAAGCTCCTCTACCACCAGGTTCCTTCTGAGCAGGAAGGCAAACGGACCCTGCGTGCGCCCACTTTCTTTGTGTCTCAAGAGGATCAATCCTTCAAGACTGAGTTCTTCCCGCCTGGTAGTGAAGCTGAGCGTCGGATCTCGTTCTTCGCGCAATCACTCTCTACCCCCATGCCCGAGCCGCTTCCTGTGGACAACATGCCTACATTCACCGTTCTGATTCCCCACTATAGCGAGAAGATCCTCCTATCCCTGCGTGAGATTATCCGCGAGGATGAGCCCTACTCTCGTGTGACGCTGCTGGAATACCTCAAACAACTTCATCCTCACGAGTGGGACTGCTTCGTCAAAGACACCAAGATTTTGGCCGA

Conidial equivalents obtained from the *FKS* gene assay.

|  | **Enumerated dose (Light microscopy)** | | **Observed conidial equivalents**  **(RT-qPCR *FKS* assay)** | | | |  |
| --- | --- | --- | --- | --- | --- | --- | --- |
|  | ***n*** | **Dose group** | **Median** | **IQR** | **Min** | **Max** | **Deviation (%)** |
| **Long-term study** | 3 | 5,000,000 (5M) | 4,894,363 | 218,126 | 4,753,181 | 5,189,433 | -2.1 |
|  | 3 | 500,000 (0.5M) | 557,797 | 35,018 | 535,401 | 605,437 | 11.6 |
|  | 3 | 50,000 (50K) | 73,589 | 1,991 | 73,589 | 77,570 | 47.2 |
|  | 3 | 5,000 (5K) | 5,766 | 2,449 | 2,221 | 7,119 | 15.3 |
| **Pilot study** | 3 | 50,000,000 (50M) | 48,530,744 | 3,100,702 | 47,407,544 | 53,608,948 | -2.9 |
|  | 3 | 5,000,000 (5M) | 5,036,382 | 150,930 | 5,006,985 | 5,308,844 | 0.7 |
|  | 3 | 500,000 (0.5M) | 470,390 | 33,308 | 462,201 | 528,817 | -5.9 |
|  | 3 | 50,000 (50K) | 73,111 | 6,621 | 64,276 | 77,519 | 46.2 |

*n* = samples tested, RT-qPCR: real-time quantitative PCR.
