## Supplementary file 6 for "New high accuracy diagnostics for avian *Aspergillus fumigatus* infection using Nanopore methylation sequencing of host cell-free DNA and machine learning prediction"

Median methylation (percentages ± IQR) of the six size classes of regions for each study and cohort, as well as the numbers of regions included for analysis.

|  |  | **Median methylation (%) across regions with varied size classes** | | | | | |
| --- | --- | --- | --- | --- | --- | --- | --- |
|  | **Status** | **1KB** | **5KB** | **10KB** | **15KB** | **20KB** | **25KB** |
| ***n* regions:** |  | 76 | 1,558 | 10,678 | 19,452 | 24,410 | 26,138 |
| Long-term study | Control | 35.9%  (± 15.6%) | 38.7%  (± 33.8%) | 44.9%  (± 28%) | 46.8%  (± 24%) | 47.9%  (± 21.3%) | 48.9%  (± 19.6%) |
|  | Infected | 36.4%  (± 16.4%) | 38.9%  (± 33.7%) | 44.9%  (± 27.6%) | 46.9%  (± 23.7%) | 48.1%  (± 21%) | 49.2%  (± 19.4%) |
| Pilot study | Control | 38%  (± 24.5%) | 38.2%  (± 39.9%) | 43.5%  (± 32.1%) | 45.5%  (± 27.7%) | 46.7%  (± 24.5%) | 47.7%  (± 22.9%) |
|  | Infected | 37.9%  (± 25.1%) | 38.9%  (± 39.6%) | 44%  (± 32.5%) | 45.8%  (± 28.1%) | 46.9%  (± 24.7%) | 47.8%  (± 22.6%) |
| Specificity cohort | Control | 34.4%  (± 18.8%) | 36.8%  (± 34.7%) | 42.2%  (± 26.3%) | 44.2%  (± 21.9%) | 45.5%  (± 19.3%) | 46.4%  (± 17.7%) |
|  | Infected | 34.6%  (± 18.6%) | 37.4%  (± 34%) | 43%  (± 26.2%) | 44.9%  (± 21.8%) | 46.2%  (± 19.3%) | 47.2%  (± 17.8%) |
| Real-world and clinical samples | Control* | 33.3%  (± 18.6%) | 33.9%  (± 39.8%) | 39%  (± 29.7%) | 40.5%  (± 24.5%) | 41.2%  (± 21.6%) | 42.1%  (± 19.6%) |
|  | Infected* | 32.4%  (± 21.7%) | 37%  (± 42.6%) | 42.9%  (± 34.4%) | 43.5%  (± 29.3%) | 43.8%  (± 25.3%) | 44.3%  (± 23%) |
| **Min –**  **Max** |  | 32.4%-  38.0% | 33.9%-  38.9% | 39.0%-  44.9% | 40.5%-  46.9% | 41.2%-  48.1% | 42.1%-  49.2% |
