## Supplementary file 7 for "New high accuracy diagnostics for avian *Aspergillus fumigatus* infection using Nanopore methylation sequencing of host cell-free DNA and machine learning prediction"

Final filtering of machine-learning (ML) models as candidates for the diagnostic tests. The three final selected tests are shown in the top three rows.

|  | **Parameters** | | | **Theoretical**  **performance** | | | | | **Observed**  **performance** | | |
| --- | --- | --- | --- | --- | --- | --- | --- | --- | --- | --- | --- |
| **Test** | **Algo.** | ***n.*** | **Size** | **F1** | **ROC-AUC** | **SE** | **SPE** | **Acc.** | **SE** | **SPE** | **Acc.** |
| **High**  **Accuracy** | **glmnet** | **83** | **10 KB** | **0.89** | **0.92** | **91.7%** | **86.1%** | **88.89%** | **100%** | **89.2%** | **92.3%** |
| **Fast** | **rf** | **22** | **10 KB** | **0.77** | **0.91** | **75.00%** | **80.60%** | **77.78%** | **93.33%** | **78.38%** | **82.69%** |
| **In-Field** | **lda** | **4** | **5 KB** | **0.67** | **0.76** | **66.70%** | **66.70%** | **66.67%** | **73.33%** | **72.97%** | **73.1%** |
| 4. | glmnet | 183 | 5KB | 0.795 | 0.89 | 80.60% | 77.80% | 79.17% | 73.33% | 78.38% | 76.92% |
| 5. | glmnet | 74 | 5KB | 0.861 | 0.94 | 86.10% | 86.10% | 86.11% | 80.00% | 67.57% | 71.15% |
| 6. | nnet | 76 | 5KB | 0.886 | 0.96 | 86.10% | 91.70% | 88.89% | 80.00% | 78.38% | 78.85% |
| 7. | nnet | 24 | 5KB | 0.778 | 0.81 | 77.80% | 77.80% | 77.78% | 86.67% | 72.97% | 76.92% |
| 8. | nnet | 78 | 5KB | 0.901 | 0.97 | 88.90% | 91.70% | 90.28% | 100.00% | 72.97% | 80.77% |
| 9. | glmnet | 39 | 5KB | 0.795 | 0.88 | 80.60% | 77.80% | 79.17% | 60.00% | 72.97% | 69.23% |
| 10. | rf | 11 | 10KB | 0.767 | 0.86 | 77.80% | 75.00% | 76.39% | 93.33% | 72.97% | 78.85% |
| 11. | glmnet | 14 | 10KB | 0.761 | 0.89 | 75.00% | 77.80% | 76.39% | 80.00% | 78.38% | 78.85% |
| 12. | NB | 21 | 10KB | 0.845 | 0.93 | 83.30% | 86.10% | 84.72% | 80.00% | 70.27% | 73.08% |
| 13. | KNN | 73 | 10KB | 0.81 | 0.91 | 88.90% | 69.40% | 79.17% | 80.00% | 81.08% | 80.77% |
| 14. | glmnet | 45 | 10KB | 0.889 | 0.93 | 88.90% | 88.90% | 88.89% | 66.67% | 83.78% | 78.85% |
| 15. | gbm | 19 | 10KB | 0.877 | 0.92 | 88.90% | 86.10% | 87.50% | 86.67% | 75.68% | 78.85% |
| 16. | NB | 22 | 10KB | 0.829 | 0.92 | 80.60% | 86.10% | 83.33% | 73.33% | 78.38% | 76.92% |
| 17. | KNN | 89 | 10KB | 0.805 | 0.9 | 86.10% | 72.20% | 79.17% | 80.00% | 86.49% | 84.62% |
| 18. | KNN | 90 | 10KB | 0.853 | 0.92 | 88.90% | 80.60% | 84.72% | 80.00% | 89.19% | 86.54% |
| 19. | KNN | 124 | 10KB | 0.831 | 0.87 | 88.90% | 75.00% | 81.94% | 66.67% | 89.19% | 82.69% |
| 20. | rf | 20 | 10KB | 0.771 | 0.85 | 75.00% | 80.60% | 77.78% | 86.67% | 78.38% | 80.77% |
| 21. | glmnet | 21 | 10KB | 0.778 | 0.87 | 77.80% | 77.80% | 77.78% | 80.00% | 78.38% | 78.85% |
| 22. | gbm | 22 | 10KB | 0.824 | 0.87 | 77.80% | 88.90% | 83.33% | 86.67% | 81.08% | 82.69% |
| 23. | rf | 17 | 10KB | 0.865 | 0.88 | 88.90% | 83.30% | 86.11% | 93.33% | 70.27% | 76.92% |
| 24. | gbm | 13 | 10KB | 0.725 | 0.78 | 69.40% | 77.80% | 73.61% | 73.33% | 78.38% | 76.92% |
| 25. | KNN | 84 | 10KB | 0.75 | 0.79 | 83.30% | 61.10% | 72.22% | 86.67% | 89.19% | 88.46% |
| 26. | KNN | 70 | 10KB | 0.769 | 0.78 | 83.30% | 66.70% | 75.00% | 86.67% | 91.89% | 90.38% |
| 27. | glmnet | 49 | 10KB | 0.849 | 0.93 | 86.10% | 83.30% | 84.72% | 66.67% | 89.19% | 82.69% |
| 28. | gbm | 50 | 10KB | 0.811 | 0.89 | 83.30% | 77.80% | 80.56% | 93.33% | 81.08% | 84.62% |
| 29. | glmnet | 72 | 10KB | 0.827 | 0.9 | 86.10% | 77.80% | 81.94% | 86.67% | 86.49% | 86.54% |
| 30. | glmnet | 155 | 10KB | 0.865 | 0.92 | 88.90% | 83.30% | 86.11% | 73.33% | 89.19% | 84.62% |
| 31. | glmnet | 74 | 10KB | 0.845 | 0.91 | 83.30% | 86.10% | 84.72% | 93.33% | 86.49% | 88.46% |
| 32. | glmnet | 77 | 10KB | 0.904 | 0.93 | 91.70% | 88.90% | 90.28% | 100.00% | 86.49% | 90.38% |
| 33. | glmnet | 10 | 15KB | 0.73 | 0.76 | 75.00% | 69.40% | 72.22% | 86.67% | 78.38% | 80.77% |
| 34. | KNN | 12 | 15KB | 0.74 | 0.76 | 75.00% | 72.20% | 73.61% | 73.33% | 78.38% | 76.92% |
| 35. | glmnet | 3 | 15KB | 0.522 | 0.54 | 50.00% | 58.30% | 54.17% | 66.67% | 81.08% | 76.92% |
| 36. | glmnet | 4 | 15KB | 0.5 | 0.56 | 50.00% | 50.00% | 50.00% | 60.00% | 91.89% | 82.69% |
| 37. | glmnet | 6 | 15KB | 0.479 | 0.55 | 47.20% | 50.00% | 48.61% | 60.00% | 94.59% | 84.62% |
| 38. | lda | 7 | 15KB | 0.611 | 0.61 | 61.10% | 61.10% | 61.11% | 80.00% | 94.59% | 90.38% |
| 39. | lda | 9 | 15KB | 0.595 | 0.59 | 61.10% | 55.60% | 58.33% | 53.33% | 94.59% | 82.69% |
| 40. | lda | 7 | 15KB | 0.514 | 0.57 | 52.80% | 47.20% | 50.00% | 33.33% | 94.59% | 76.92% |
| 41. | rf | 91 | 15KB | 0.72 | 0.83 | 75.00% | 66.70% | 70.83% | 80.00% | 86.49% | 84.62% |
| 42. | gbm | 5 | 20KB | 0.638 | 0.67 | 61.10% | 69.40% | 65.28% | 33.33% | 81.08% | 67.31% |
| 43. | lda | 6 | 20KB | 0.56 | 0.56 | 58.30% | 50.00% | 54.17% | 86.67% | 78.38% | 80.77% |
| 44. | lda | 7 | 20KB | 0.568 | 0.51 | 58.30% | 52.80% | 55.56% | 80.00% | 81.08% | 80.77% |
| 45. | rf | 50 | 20KB | 0.72 | 0.77 | 75.00% | 66.70% | 70.83% | 86.67% | 89.19% | 88.46% |
| 46. | rf | 54 | 20KB | 0.722 | 0.78 | 72.20% | 72.20% | 72.22% | 73.33% | 91.89% | 86.54% |
| 47. | lda | 7 | 25KB | 0.712 | 0.81 | 72.20% | 69.40% | 70.83% | 60.00% | 78.38% | 73.08% |
| 48. | nnet | 158 | 25KB | 0.986 | 1 | 100.00% | 97.20% | 98.61% | 86.67% | 83.78% | 84.62% |
| 49. | gbm | 9 | 25KB | 0.648 | 0.63 | 63.90% | 66.70% | 65.28% | 46.67% | 89.19% | 76.92% |
