## Supplementary file 8 for "New high accuracy diagnostics for avian *Aspergillus fumigatus* infection using Nanopore methylation sequencing of host cell-free DNA and machine learning prediction"

Differentially methylated cell-free DNA regions (DMR) as diagnostic markers included in each of the three tests.

**High accuracy test**

| **ID** | **chr** | **start** | **end** | **pvalue** | **qvalue** | **meth.diff** |
| --- | --- | --- | --- | --- | --- | --- |
| HAC_01 | 16 | 520000 | 530000 | 0 | 0 | 5.41 |
| HAC_02 | 16 | 770000 | 780000 | 0 | 0 | 6.05 |
| HAC_03 | 16 | 1630000 | 1640000 | 0 | 0 | 5.11 |
| HAC_04 | 16 | 480000 | 490000 | 7.77E-216 | 9.22E-213 | 5.57 |
| HAC_05 | 16 | 1030000 | 1040000 | 1.19E-164 | 9.79E-163 | 5.10 |
| HAC_06 | 16 | 1160000 | 1170000 | 2.68E-81 | 1.51E-78 | 6.80 |
| HAC_07 | 16 | 820000 | 830000 | 7.37E-72 | 3.93E-69 | 5.36 |
| HAC_08 | 6 | 20800000 | 20810000 | 1.02E-43 | 4.55E-41 | 9.73 |
| HAC_09 | 16 | 400000 | 410000 | 1.01E-42 | 4.32E-40 | 7.83 |
| HAC_10 | 1 | 1.96E+08 | 1.96E+08 | 1.10E-30 | 4.06E-28 | 6.17 |
| HAC_11 | 16 | 1130000 | 1140000 | 4.85E-24 | 1.62E-21 | 5.38 |
| HAC_12 | 27 | 3460000 | 3470000 | 8.00E-23 | 2.59E-18 | 8.31 |
| HAC_13 | 25 | 2000000 | 2010000 | 1.24E-14 | 3.79E-12 | 6.14 |
| HAC_14 | 5 | 26920000 | 26930000 | 8.32E-14 | 2.47E-11 | 7.12 |
| HAC_15 | 5 | 54400000 | 54410000 | 1.18E-13 | 3.42E-11 | 7.67 |
| HAC_16 | 17 | 7240000 | 7250000 | 1.78E-13 | 5.00E-11 | 7.66 |
| HAC_17 | Z | 27650000 | 27660000 | 2.45E-13 | 6.71E-13 | -5.98 |
| HAC_18 | 25 | 1890000 | 1900000 | 5.36E-11 | 1.36E-08 | 5.15 |
| HAC_19 | 19 | 5980000 | 5990000 | 7.50E-11 | 1.86E-08 | 5.58 |
| HAC_20 | 17 | 1320000 | 1330000 | 1.15E-09 | 2.80E-07 | 5.24 |
| HAC_21 | 39 | 80000 | 90000 | 2.92E-09 | 6.93E-07 | 6.10 |
| HAC_22 | 17 | 1780000 | 1790000 | 4.70E-09 | 1.09E-06 | 8.06 |
| HAC_23 | 4 | 28780000 | 28790000 | 9.90E-09 | 2.25E-06 | 6.71 |
| HAC_24 | Z | 27780000 | 27790000 | 1.04E-08 | 2.31E-06 | -5.31 |
| HAC_25 | 11 | 6290000 | 6300000 | 1.58E-08 | 3.44E-06 | 7.51 |
| HAC_26 | 10 | 8680000 | 8690000 | 6.75E-08 | 1.41E-05 | 6.86 |
| HAC_27 | 28 | 3320000 | 3330000 | 7.51E-09 | 1.54E-05 | 5.49 |
| HAC_28 | 6 | 26610000 | 26620000 | 6.19E-07 | 1.22E-04 | 8.05 |
| HAC_29 | 19 | 7680000 | 7690000 | 6.18E-06 | 1.20E-03 | 5.34 |
| HAC_30 | 3 | 42630000 | 42640000 | 2.90E-05 | 5.43E-03 | 5.44 |
| HAC_31 | 15 | 7520000 | 7530000 | 6.50E-06 | 1.18E-02 | -4.33 |
| HAC_32 | 31 | 80000 | 90000 | 7.45E-05 | 1.33E-02 | -7.82 |
| HAC_33 | Z | 27670000 | 27680000 | 9.28E-05 | 1.60E-02 | -5.34 |
| HAC_34 | 5 | 21200000 | 21210000 | 2.23E-05 | 3.61E-02 | 5.21 |
| HAC_35 | 2 | 30910000 | 30920000 | 5.71E-04 | 8.96E-02 | 5.46 |
| HAC_36 | 1 | 84040000 | 84050000 | 1.24E-03 | 1.81E-02 | 5.70 |
| HAC_37 | 5 | 38650000 | 38660000 | 3.09E-03 | 4.23E-01 | 6.06 |
| HAC_38 | 1 | 80580000 | 80590000 | 9.11E-03 | 1.20E+00 | 5.09 |
| HAC_39 | 5 | 53760000 | 53770000 | 1.49E-02 | 1.94E+00 | 5.71 |
| HAC_40 | 14 | 1120000 | 1130000 | 3.89E-02 | 4.83E+00 | 5.55 |
| HAC_41 | 12 | 2550000 | 2560000 | 4.73E-02 | 5.81E+00 | 5.29 |
| HAC_42 | 2 | 1.28E+08 | 1.28E+08 | 1.77E-01 | 2.10E+01 | 5.48 |
| HAC_43 | 14 | 990000 | 1000000 | 1.86E-01 | 2.16E+01 | 6.24 |
| HAC_44 | Z | 27680000 | 27690000 | 2.30E-01 | 2.64E+01 | -4.88 |
| HAC_45 | 14 | 15180000 | 15190000 | 5.83E-01 | 6.16E+01 | 6.63 |
| HAC_46 | 4 | 14170000 | 14180000 | 8.51E-01 | 8.65E+01 | 5.35 |
| HAC_47 | Z | 27660000 | 27670000 | 1.28E+00 | 1.28E+02 | -4.02 |
| HAC_48 | 19 | 3900000 | 3910000 | 1.37E+00 | 1.35E+02 | 5.28 |
| HAC_49 | 7 | 2380000 | 2390000 | 1.41E+00 | 1.38E+02 | 5.26 |
| HAC_50 | 1 | 1.32E+08 | 1.32E+08 | 3.59E+00 | 3.45E+02 | -5.05 |
| HAC_51 | Z | 27770000 | 27780000 | 3.70E+00 | 3.53E+02 | -4.25 |
| HAC_52 | 11 | 17740000 | 17750000 | 3.88E+00 | 3.64E+02 | 5.83 |
| HAC_53 | 10 | 8910000 | 8920000 | 5.97E+00 | 5.31E+02 | 5.49 |
| HAC_54 | 5 | 23930000 | 23940000 | 7.80E+00 | 6.83E+02 | 5.66 |
| HAC_55 | 25 | 1090000 | 1100000 | 1.39E+01 | 1.19E+02 | -4.16 |
| HAC_56 | 2 | 4060000 | 4070000 | 1.63E+01 | 1.39E+03 | -5.15 |
| HAC_57 | 2 | 69070000 | 69080000 | 1.99E+01 | 1.64E+03 | 6.18 |
| HAC_58 | 10 | 1860000 | 1870000 | 2.72E+01 | 2.24E+03 | 5.02 |
| HAC_59 | 19 | 8670000 | 8680000 | 3.36E+01 | 2.71E+03 | 6.49 |
| HAC_60 | 1 | 38030000 | 38040000 | 4.00E+01 | 3.10E+03 | 5.56 |
| HAC_61 | 3 | 58890000 | 58900000 | 3.90E+01 | 3.10E+03 | 5.23 |
| HAC_62 | 10 | 1200000 | 1210000 | 4.01E+01 | 3.10E+03 | -4.00 |
| HAC_63 | 2 | 780000 | 790000 | 4.86E+01 | 3.66E+02 | 5.15 |
| HAC_64 | 26 | 4340000 | 4350000 | 7.01E+01 | 5.13E+03 | 6.16 |
| HAC_65 | 12 | 1940000 | 1950000 | 8.24E+01 | 5.98E+03 | -2.82 |
| HAC_66 | Z | 27640000 | 27650000 | 1.08E+02 | 7.63E+03 | -4.33 |
| HAC_67 | 6 | 9750000 | 9760000 | 1.41E+02 | 9.68E+03 | 5.82 |
| HAC_68 | 5 | 21100000 | 21110000 | 2.13E+02 | 1.42E+04 | 5.03 |
| HAC_69 | 2 | 30680000 | 30690000 | 2.38E+02 | 1.57E+04 | 5.26 |
| HAC_70 | 4 | 59420000 | 59430000 | 3.44E+02 | 2.24E+03 | 6.16 |
| HAC_71 | 22 | 160000 | 170000 | 6.10E+02 | 3.77E+04 | 5.47 |
| HAC_72 | 7 | 15890000 | 15900000 | 1.78E+03 | 1.04E+05 | -3.28 |
| HAC_73 | 39 | 120000 | 130000 | 1.84E+03 | 1.06E+05 | -3.61 |
| HAC_74 | 5 | 43620000 | 43630000 | 2.77E+03 | 1.53E+05 | 5.81 |
| HAC_75 | 9 | 2270000 | 2280000 | 3.17E+03 | 1.72E+05 | -3.81 |
| HAC_76 | 2 | 1.03E+08 | 1.03E+08 | 3.49E+03 | 1.86E+05 | 5.10 |
| HAC_77 | 1 | 71380000 | 71390000 | 1.58E+04 | 7.43E+05 | -4.90 |
| HAC_78 | 9 | 14970000 | 14980000 | 1.97E+04 | 8.92E+05 | 5.72 |
| HAC_79 | 15 | 5240000 | 5250000 | 1.96E+04 | 8.92E+05 | -5.25 |
| HAC_80 | 1 | 82260000 | 82270000 | 1.99E+04 | 8.95E+05 | 5.12 |
| HAC_81 | 3 | 66260000 | 66270000 | 5.75E+04 | 2.35E+05 | 6.39 |
| HAC_82 | 4 | 44900000 | 44910000 | 6.08E+03 | 2.46E+06 | 5.06 |
| HAC_83 | 4 | 73190000 | 73200000 | 6.28E+04 | 2.51E+06 | 5.12 |

**Fast test**

| **ID** | **chr** | **start** | **end** | **pvalue** | **qvalue** | **meth.diff** |
| --- | --- | --- | --- | --- | --- | --- |
| FAST_01 | 16 | 480000 | 490000 | 3.24E-232 | 4.94E-229 | 5.71 |
| FAST_02 | 16 | 1160000 | 1170000 | 3.21E-64 | 1.72E-61 | 6.07 |
| FAST_03 | 16 | 400000 | 410000 | 1.76E-33 | 6.98E-31 | 7.11 |
| FAST_04 | 6 | 20800000 | 20810000 | 1.74E-20 | 6.63E-18 | 7.35 |
| FAST_05 | 1 | 1.96E+08 | 1.96E+08 | 3.53E-19 | 1.26E-15 | 5.17 |
| FAST_06 | 1 | 71380000 | 71390000 | 3.08E-16 | 9.97E-14 | -8.57 |
| FAST_07 | 27 | 3460000 | 3470000 | 6.76E-09 | 1.90E-07 | 6.53 |
| FAST_08 | 28 | 3320000 | 3330000 | 1.66E-08 | 4.55E-06 | 5.53 |
| FAST_09 | 25 | 2000000 | 2010000 | 3.75E-08 | 1.00E-05 | 5.29 |
| FAST_10 | 31 | 80000 | 90000 | 3.89E-05 | 9.04E-03 | -7.79 |
| FAST_11 | 20 | 1560000 | 1570000 | 5.21E-03 | 1.03E+00 | 6.58 |
| FAST_12 | 4 | 42940000 | 42950000 | 1.22E-02 | 2.36E+00 | 5.14 |
| FAST_13 | 5 | 54400000 | 54410000 | 5.70E-02 | 1.05E+01 | 5.67 |
| FAST_14 | 10 | 8910000 | 8920000 | 1.23E+00 | 2.06E+02 | 5.59 |
| FAST_15 | 17 | 1780000 | 1790000 | 3.10E+00 | 5.01E+02 | 6.13 |
| FAST_16 | 25 | 130000 | 140000 | 1.03E+01 | 1.30E+04 | 5.24 |
| FAST_17 | 4 | 73190000 | 73200000 | 3.98E+02 | 4.67E+04 | 5.71 |
| FAST_18 | Z | 1700000 | 1710000 | 5.19E+03 | 5.09E+04 | 5.01 |
| FAST_19 | 1 | 82260000 | 82270000 | 1.35E+04 | 1.22E+06 | 5.11 |
| FAST_20 | 6 | 26610000 | 26620000 | 2.71E+05 | 1.91E+07 | 5.04 |
| FAST_21 | 17 | 6150000 | 6160000 | 2.76E+05 | 1.92E+07 | 5.40 |
| FAST_22 | 3 | 66260000 | 66270000 | 1.29E+07 | 5.49E+08 | 5.41 |

**In situ test**

| ID | chr | start | end | pvalue | qvalue | meth.diff |
| --- | --- | --- | --- | --- | --- | --- |
| FIELD_01 | 16 | 770000 | 775000 | 4.46E-100 | 3.48E-99 | 8.85 |
| FIELD_02 | 5 | 21200000 | 21205000 | 1.48E-06 | 4.34E-06 | 7.23 |
| FIELD_03 | 12 | 1605000 | 1610000 | 5.08E-02 | 1.36E+00 | 7.56 |
| FIELD_04 | Z | 27670000 | 27675000 | 1.22E-01 | 3.21E+00 | -6.38 |
