## Supplementary file 9 for "New high accuracy diagnostics for avian *Aspergillus fumigatus* infection using Nanopore methylation sequencing of host cell-free DNA and machine learning prediction"

**Median methylation values based on infection status with statistical comparison using Wilcoxon.**

|  |  | **Control** | | **Infected** | |  |
| --- | --- | --- | --- | --- | --- | --- |
| **Week** | **Test** | **Median methylation** | **IQR** | **Median methylation** | **IQR** | **P-value** |
| Week 2 | High accuracy test | 62.1% | 26.2% | 62.6% | 28.3% | 0.67 |
|  | Fast test | 55.4% | 24.7% | 56.7% | 25.1% | 0.24 |
|  | In situ test | 60.2% | 25.5% | 63.8% | 20.5% | 0.19 |
| Week 4 | High accuracy test | 57.9% | 26.4% | 59.5% | 26.8% | 0.11 |
|  | Fast test | 51.4% | 22.9% | 53.3% | 23.8% | 0.06 ‘ |
|  | In situ test | 56.9% | 24.2% | 57.4% | 32.7% | 0.45 |
| Week 5 | **High accuracy test** | **58.6%** | **25.7%** | **62.9%** | **24.2%** | **0.00 ***** |
|  | **Fast test** | **52.3%** | **21.5%** | **57.5%** | **23.3%** | **0.00 ***** |
|  | In situ test | 62.8% | 23.5% | 66.7% | 25.7% | 0.21 |

IQR: interquartile range.

|  |  | **Control** | | **Infected** | |  |
| --- | --- | --- | --- | --- | --- | --- |
| **Cohort** | **Test** | **Median methylation** | **IQR** | **Median methylation** | **IQR** | **P-value** |
| Pilot Study | **High accuracy test** | **58.0%** | **37.2%** | **63.2%** | **37.8%** | **0.00 ***** |
|  | Fast test | 54.0% | 33.2% | 56.3% | 32.2% | 0.07 ‘ |
|  | In situ test | 59.0% | 38.4% | 71.8% | 24.6% | 0.12 |
| Specificity cohort –  *G. anatis* | High accuracy test | **49.2%** | **29.9%** | **54.1%** | **30.1%** | **0.05 *** |
|  | Fast test | 44.7% | 21.9% | 48.6% | 26.8% | 0.46 |
|  | In situ test | 52.6% | 26.9% | 58.0% | 24.2% | 0.26 |
| Specificity cohort –  *E. coli* | High accuracy test | 52.1% | 27.9% | 53.7% | 29.0% | 0.45 |
|  | Fast test | 49.8% | 28.2% | 52.8% | 27.7% | 0.25 |
|  | In situ test | 46.1% | 34.1% | 45.8% | 36.0% | 0.70 |
| Clinical samples | **High accuracy test** | **52.8%** | **33.0%** | **64.0%** | **32.7%** | **0.00 ***** |
|  | **Fast test** | **42.7%** | **27.1%** | **60.6%** | **31.2%** | **0.00 ***** |
|  | In situ test | 71.1% | 18.8% | 73.4% | 19.2% | 0.86 |

IQR: interquartile range.
