## Supplementary file 10 for "New high accuracy diagnostics for avian *Aspergillus fumigatus* infection using Nanopore methylation sequencing of host cell-free DNA and machine learning prediction"

Post-mortem scoring sheet with scoring grading definitions for each individual parameter.

| **Group:** | | **Animal ID:** | **Score** | | | | |
| --- | --- | --- | --- | --- | --- | --- | --- |
|  |  |  | 0 | 1 | 2 | 3 | 4 |
| **Peritoneum** | **P1** | Inflammatory reaction |  |  |  |  |  |
|  | **P2** | Amount of exudate |  |  |  |  |  |
|  | **P3** | Type of exudate |  |  |  |  |  |
|  | **P4** | Transparency of peritoneum |  |  |  |  |  |
| **(Left) Airsac** | **A1** | Transparency of airsac |  |  |  |  |  |
|  | **A2** | Amount of exudate |  |  |  |  |  |
| **(Left) Lung**    Left lung weight (g): | **L1** | Amount of exudate |  |  |  |  |  |
|  | **L2** | Congestion |  |  |  |  |  |
|  | **L3** | Lung lesions (oedema) |  |  |  |  |  |
|  | **L4** | Lung lesions (size) |  |  |  |  |  |
|  | **L5** | Granulomas |  |  |  |  |  |
| **Spleen**      Weight (g):____________ | **Spl1** | Proliferation |  |  |  |  |  |
|  | **Spl2** | Enlarged  Size in cm: lenght:_____      width______ |  |  |  |  |  |
| **Other** |  | Liver |  |  |  |  |  |
|  |  | Trachea |  |  |  |  |  |

**Scoring definitions:**

| **P1** | Inflammatory reaction | 0: None  1: Local at the entrance  2: Around the ovary  3: In the omentum  4: All peritoneum |
| --- | --- | --- |
| **P2** | Amount of exudate | 0: None  1: Sparse  2: Some  3: Abundant |
| **P3** | Type of exudate | 0: None  1: Aqueous  2: Lump of pus  3: Flakes  4: Confluent |
| **P4** | Transparency of peritoneum | 0: Clear  1: Unclear  2: Cloudy  3: Milky  4: Opaque |
| **A1** | Transparency of airsac | 0: Clear  1: Unclear  2: Cloudy  3: Milky  4: Opaque |
| **A2** | Amount of exudate | 0: None  1: Sparse  2: Some  3: Abundant |
| **L1** | Amount of exudate | 0: None  1: Sparse  2: Some  3: Abundant |
| **L2** | Congestion | 0: None  1: Sparse (<25% of lung tissue)  2: Pronounced (>25% of lung tissue) |
| **L3** | Lung lesions (oedema) | 0: None.  1: Slight oedema of the alveolar walls.  2: Moderate oedematous thickening of alveolar walls with occasional alveoli containing coagulated oedema fluids.  3: Extensive occurrence of alveolar and interstitial oedema. |
| **L4** | Lung lesions (size) | 0: no lesions  1: small lesions between 1 cm2 and 2 cm2  2: moderate-size lesions between 3 cm2 and 4 cm2  3: extensive lesions with more than 4 cm2  and covering almost all the area of the lungs |
| **L5** | Granulomas | 0: None  1: Few small white caseous/spherical nodules  2: Numerous small white caseous/spherical nodules  3: Many caseous lesions/nodules and green-gray mold due to sporulation |
| **Spl1** | Lymphoid reaction (Proliferation) | 0: None  1: Little proliferation  2: Distinct proliferation |
| **Spl2** | Enlargement | 0: None  1: Less than double size  2: More than double size  3: Atrophic/depleted |
